# LipIDens: Simulation assisted interpretation of lipid densities in cryo-EM structures of membrane proteins

**DOI:** 10.1101/2022.06.30.498233

**Authors:** T. Bertie Ansell, Wanling Song, Claire E. Coupland, Loic Carrique, Robin A. Corey, Anna L. Duncan, C. Keith Cassidy, Maxwell M. G. Geurts, Tim Rasmussen, Andrew B. Ward, Christian Siebold, Phillip J. Stansfeld, Mark S. P. Sansom

**Affiliations:** Department of Biochemistry, University of Oxford, University of Oxford, South Parks Road, Oxford, OX1 3QU, UK; Division of Structural Biology, Wellcome Centre for Human Genetics, University of Oxford, Roosevelt Drive, Oxford, OX3 7BN, UK; Biocenter and Rudolf-Virchow-Zentrum, Universität Würzburg, Haus D15, Josef-Schneider-Str. 2, 97080 Würzburg, Germany; Department of Integrative Structural and Computational Biology, The Scripps Research Institute, La Jolla, California 92037, USA; School of Life Sciences & Department of Chemistry, University of Warwick, Coventry, CV4 7AL, UK; Odyssey Therapeutics, London, UK

**Keywords:** Simulation, lipid, molecular dynamics, cryo-EM, density, structure, membrane protein

## Abstract

Cryo-electron microscopy (cryo-EM) enables the determination of membrane protein structures in native-like environments. Characterising how membrane proteins interact with the surrounding membrane lipid environment is assisted by resolution of lipid-like densities visible in cryo-EM maps. Nevertheless, establishing the molecular identity of putative lipid and/or detergent densities remains challenging. Here we present LipIDens, a pipeline for molecular dynamics (MD) simulation-assisted interpretation of lipid and lipid-like densities in cryo-EM structures. The pipeline integrates the implementation and analysis of multi-scale MD simulations for identification, ranking and refinement of lipid binding poses which superpose onto cryo-EM map densities. Thus, LipIDens enables direct integration of experimental and computational structural approaches to facilitate the interpretation of lipid-like cryo-EM densities and to reveal the molecular identities of protein-lipid interactions within a bilayer environment. The LipIDens code is open-source and embedded within a notebook format to assist automation and usability.

## Introduction

Recent methodological advances in cryo-electron microscopy (cryo-EM) have transformed our understanding of membrane protein structure and function^1,2^. As these methods develop and enable determination of higher resolution membrane protein structures^3–8^, additional non-protein lipid-like densities are increasingly resolved surrounding protein transmembrane domains (TMDs)^9–11^. These additional densities are generally considered to correspond to bound lipid or detergent molecules. However, determining the chemical identity of putative lipid/detergent densities from cryo-EM maps is challenging^4,11,12^. As such, assignment and discussion of lipid-like densities is often tentative, complicating subsequent interpretation of how bound lipids and the bilayer environment may modulate membrane protein function.

Molecular dynamics (MD) simulations enable exploration of the lipid environment surrounding membrane proteins and have been readily applied to characterise lipid binding sites on diverse family members including G-protein coupled receptors, solute transporters, and ion channels^13–17^. In such simulations, the identity of a lipid bound at a site is known precisely. However, accompanying experimental validation of the lipid species at a predicted binding site in a native cell membrane is often absent or at best difficult to obtain. Thorough exploration of the surrounding membrane environment requires simulation timescales that are sufficient to sample multiple lipid binding/unbinding events across the TMD^14,18^. This is readily enabled through use of coarse-grained (CG) and atomistic simulations which have been used to successfully predict lipid binding sites subsequently validated via experimental structural and biophysical methods^19–21^. Thus, there is a clear complementarity between MD simulations and structure determination by cryo-EM for identification and characterisation of protein-lipid interactions. However, automated and objective protocols for exploiting this complementarity have yet to be made available.

Recent advances in software development have sought to standardise methods for determining protein-lipid interactions from simulations^22–24^. We recently developed the protein-lipid analysis toolkit, PyLipID^24^, which uses a community analysis-based approach to identify lipid binding sites and to characterise the kinetics of the binding sites and their associated residues (see ^24^ for details). PyLipID is a powerful standalone tool, however the interpretation of PyLipID outputs is dependent on a) the setup of the input MD simulations and b) effective post-processing and assessment of PyLipID outputs. Additional atomistic simulations may also be needed to refine observed lipid interactions. This therefore prompted the development of LipIDens, an integrated pipeline for assisted interpretation of lipid-like cryo-EM densities using multi-scale MD simulations. Outputs of the pipeline include representative lipid binding poses at sites where corresponding lipid-like densities are observed, including quantitative assessment of how well these match using Q scores^25^. Importantly, LipIDens can be used to rank the binding site kinetics of different lipid species at a binding site, and therefore aid identification of the most likely lipid accounting for observed structural densities. These can be used to refine lipid binding poses during model building in cryo-EM and assist structural interpretation. Thus, we provide a formalised pipeline interlacing simulation methodologies with structural characterisation of lipid-like densities; a frequently encountered and nuanced challenge in membrane protein structural biology.

## Results

### The LipIDens pipeline

An overview of the LipIDens pipeline is shown in Fig. 1. The LipIDens pipeline can be broken into multiple sections corresponding to: a) structure processing; b) setting up and performing CG simulations; c) testing PyLipID cut-offs; d) selecting PyLipID input parameters and running PyLipID analysis; e) screening PyLipID data; f) comparing lipid poses with cryo-EM densities; g) ranking site lipids; and h) lipid pose refinement using atomistic simulations. Pipeline steps are integrated into a computational notebook to assist automation (https://github.com/TBGAnsell/LipIDens/blob/main/LipIDens.ipynb) and detailed within the accompanying procedure. A standalone python file also permits modular implementation of LipIDens stages (https://github.com/TBGAnsell/LipIDens/blob/main/lipidens_master_run.py).

**Figure 1:**
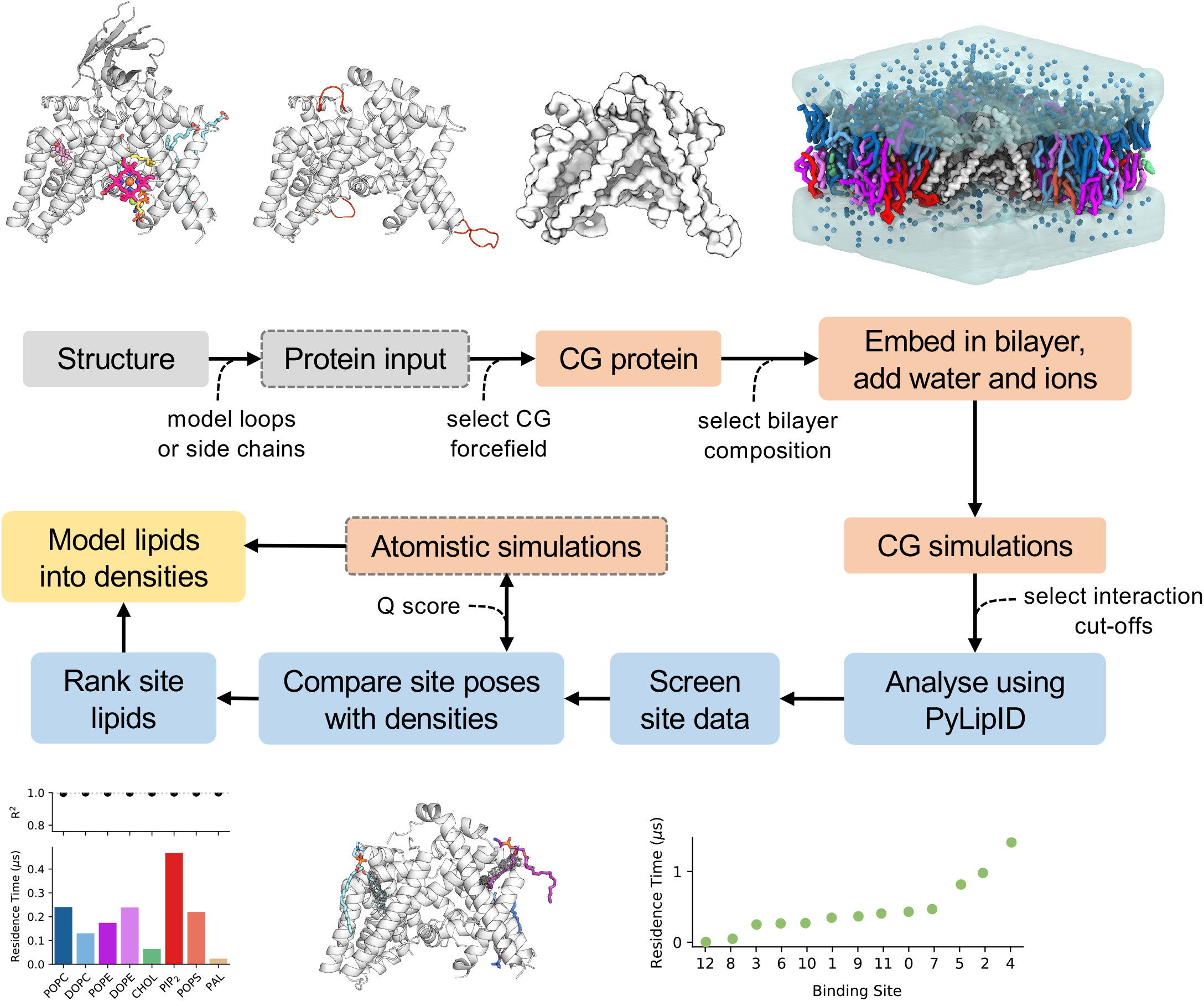
The LipIDens pipeline for characterising lipid densities using simulations. A workflow for LipIDens assisted interpretation of lipid densities using simulations, applied to Hedgehog acyltransferase (HHAT, PDBid: 7Q1U)^27^ enzyme as an example. Steps involving structure processing (grey), setup and performing MD simulations (orange), analysis of lipid sites/densities (blue) and modelling (yellow) are indicated. Optional steps are boxed by grey dashed lines. A protein structure is used as input and, if required, missing peptide linkages and/or residue sidechains are amended in the input structure. Superfluous protein components e.g. nanobodies/ligands are removed. The protein is converted to coarse-grained (CG) resolution and embedded in a selected membrane environment which is solvated using water and ions. CG simulations are performed and analysed using the lipid interaction analysis toolkit PyLipID^24^. Lipid binding sites and poses identified by PyLipID are processed, ranked and compared to densities in the cryo-EM map to assist interpretation of putative lipid densities in the structure.

### Applications

The pipeline can be used to:

- Assess whether adjacent tail-like densities observed in a cryo-EM map are likely to belong to the same or different binding sites.
- Assess how the properties of a site might favour preferential binding of one lipid type over another by examining the relative residence times of distinct lipid species binding to the same site. This can aid interpretation of structurefunction relationships.
- Obtain a more complete picture of lipid interactions within the context of a native-like membrane. This may reveal transient lipid interaction sites which are less likely to survive the purification strategies used in cryo-EM, as well as highlight the importance of lipid-lipid interactions, such as cholesterol stacking^26^.
- Quantify the kinetics of lipid binding to different sites or of multiple lipids binding to the same site. This can be used infer which sites may be more important in a biological context.
- Assess differences in lipid binding properties compared with related detergent densities.
- Check whether sterol derivates such as cholesterol-hemisuccinate, commonly used as detergents in protein purification, bind in a similar location to cholesterol in simulations. This can aid differentiation of sterol-like *vs*. phospholipid-like densities.
- Assess the relative contribution of a lipid headgroup *vs*. hydrophobic acyl tail to the interactions at a binding site.
- Enable iterative simulation and model building cycles in cryo-EM.

### Pipeline implementation

We applied the LipIDens pipeline to a recent ~2.7 Å cryo-EM structure of the ER resident enzyme, Hedgehog acyltransferase (HHAT)^27^ (Fig. 1-4). The structure of HHAT reveals several lipid-like densities, evenly distributed around the TMD, including two densities which protrude into the enzyme core. LipIDens was used to establish CG simulations of HHAT in a native-like bilayer environment. After performing CG simulations, we used LipIDens to screen dual cut-off interaction schemes for subsequent PyLipID analysis, exemplified for phosphatidylinositol 4,5-bisphosphate (PIP_2_) (Fig. 2a, Extended Data Fig. 1). During cut-off screening the minimum distances of each interacting PIP_2_ to a residue are calculated (Fig. 2a, Extended Data Fig. 1a-d) in addition to exhaustive screening of interactions over multiple cut-off pairs (Extended Data Fig. 1e-g). The selected lower cut-off (0.475 nm) corresponds to the first peak in the probability distribution plot (Fig. 2a) and the cut-off at which there is an increase in interaction durations, computed binding sites and residues comprising each site compared with smaller lower cut-off values (Extended Data Fig. 1e-g). The upper cutoff captures the first interaction shell in the probability density distribution (0.7 nm), corresponding approximately to the position of the minimum between the first and second peaks (Fig. 2a).

**Figure 2:**
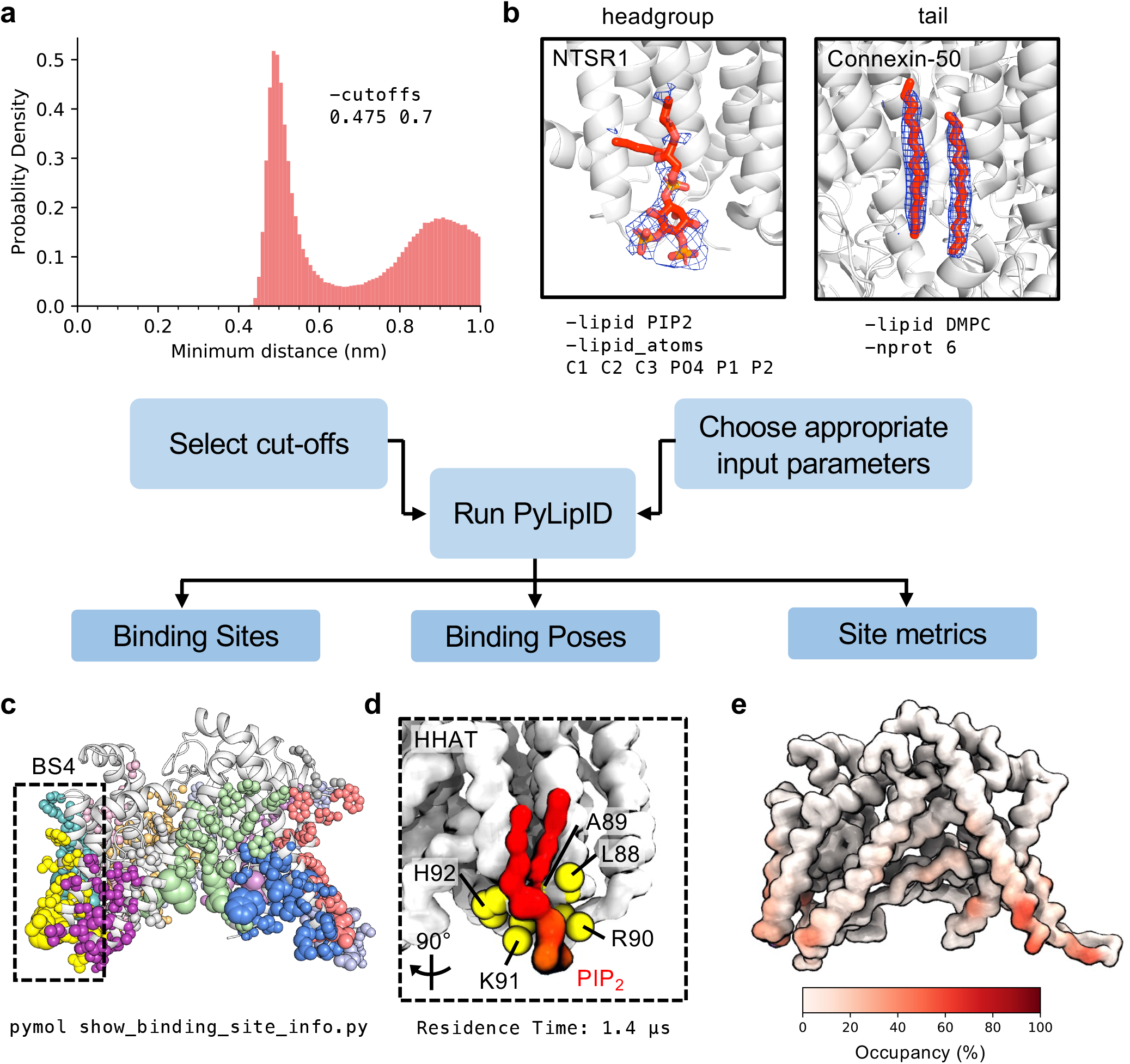
Analysing simulations using PyLipID. **a)** The upper and lower distance cut-offs used to define lipid contacts with a protein are selected from a probability distribution of the lipid of interest around the protein; exemplified here for PIP_2_ binding to HHAT. **b)** The user can tune appropriate inputs for the lipid interaction analysis using PyLipID^24^. For example, if only headgroup density is visible the user may limit the selection to lipid headgroup atoms. This is exemplified for a PIP_2_ (red sticks) binding on the neurotensin receptor (NTSR1, white cartoon). Density modelled as the PIP_2_ headgroup is shown as blue mesh (PDBid: 6UP7)^47^. Alternatively if tail density is visible the user may choose to analyse the whole lipid, as exemplified for densities (blue mesh) visible surrounding the Connexin-50 gap junction channel (PDBid: 7JJP, white cartoon)^5^. Analysis can also be averaged over homo-multimeric proteins to enhance sampling of lipid interactions. **c-e)** Example outputs from PyLipID analysis of PIP_2_ binding to HHAT from 10 x 15 μs CG simulations. A 0.475/0.7 nm dual cut-off was used to analyse interactions with the whole PIP_2_ lipid. **c)** PIP_2_ binding sites mapped onto the structure of HHAT. Binding sites are coloured individually and residues comprising each site are shown as spheres, scaled by residence time. The binding site (BS) with the longest residence time (BS4) is boxed. **d)** CG representation of the highest ranked lipid binding pose for PIP_2_ (red) at BS4. HHAT is shown in white and the top 5 residues with highest residence times within BS4 are shown as yellow spheres. **e)** PIP_2_ interactions occupancies mapped onto the structure of HHAT, coloured from low (white) to high (red).

Next, PyLipID implements this dual interaction distance cut-off (i.e. 0.475/0.7 nm) to robustly capture lipid interactions and account for transient deviations in their position due to Brownian motion^28^. Input lipid atoms may also be tuned to match structural densities (if required) i.e., by including only headgroup atoms or averaging over protein subunits (Fig. 2b). Lipid interaction durations are used to obtain the normalised survival time correlation function (hereafter survival function) of interactions. A dissociation rate constant (*k_off_*) for lipid interactions with a residue is obtained by biexponential curve fitting of the interaction survival function alongside bootstrapping to the same data. PyLipID can also identify binding sites by grouping residues which simultaneously interact with the same bound lipid molecule, based upon a community analysis approach^29,30^, as shown for PIP_2_ sites mapped onto the HHAT structure using an automatically-generated PyMOL script (Fig. 2c). Kinetic parameters are then obtained for each predicted binding site. Representative lipid binding poses at a site are obtained by empirical scoring of lipid binding poses against the simulation-derived lipid density within the site. Here the representative PIP_2_ pose at the site with longest residence time (BS4) is shown (Fig. 2d). In addition, lipid interaction occupancies are calculated as the percentage of frames where lipid is bound compared to the total number of frames on a per residue or site basis (Fig. 2e). The methodological underpinnings of PyLipID are described extensively elsewhere^24^ and have been applied to a number of recent examples^31–33^.

After calculation of lipid binding sites and their kinetics, the LipIDens pipeline ranks site outputs for inspection of site quality. Site occupancies, residence times and surface areas are ranked from lowest to highest or closest to 0 for Δ*k_off_* (defined as the difference between *k_off_* calculated by curve-fitting and via bootstrapping the same data) (Fig. 3a). This plot can be used to inspect the quality of calculated binding sites. Typically, a good site has a Δ*k_off_* between ± 1 μs. For example, for HHAT, binding site 12 is ranked last by all metrics whereas binding site 4 (Fig 3a, Fig. 2c-d) has the longest predicted residence time and occupancy and a small Δ*k_off_* indicating good agreement between *k_off_* values calculated from the survival function (Fig. 3b). Poorly fitted sites, indicated by large Δ*k_off_* values and/or sparse interaction duration plots (Fig. 3c) should be excluded in subsequent stages of the pipeline. Thus, the LipIDens pipeline employs automated steps to guide users through structure and simulation processing and assess the quality of interaction outputs.

**Figure 3:**
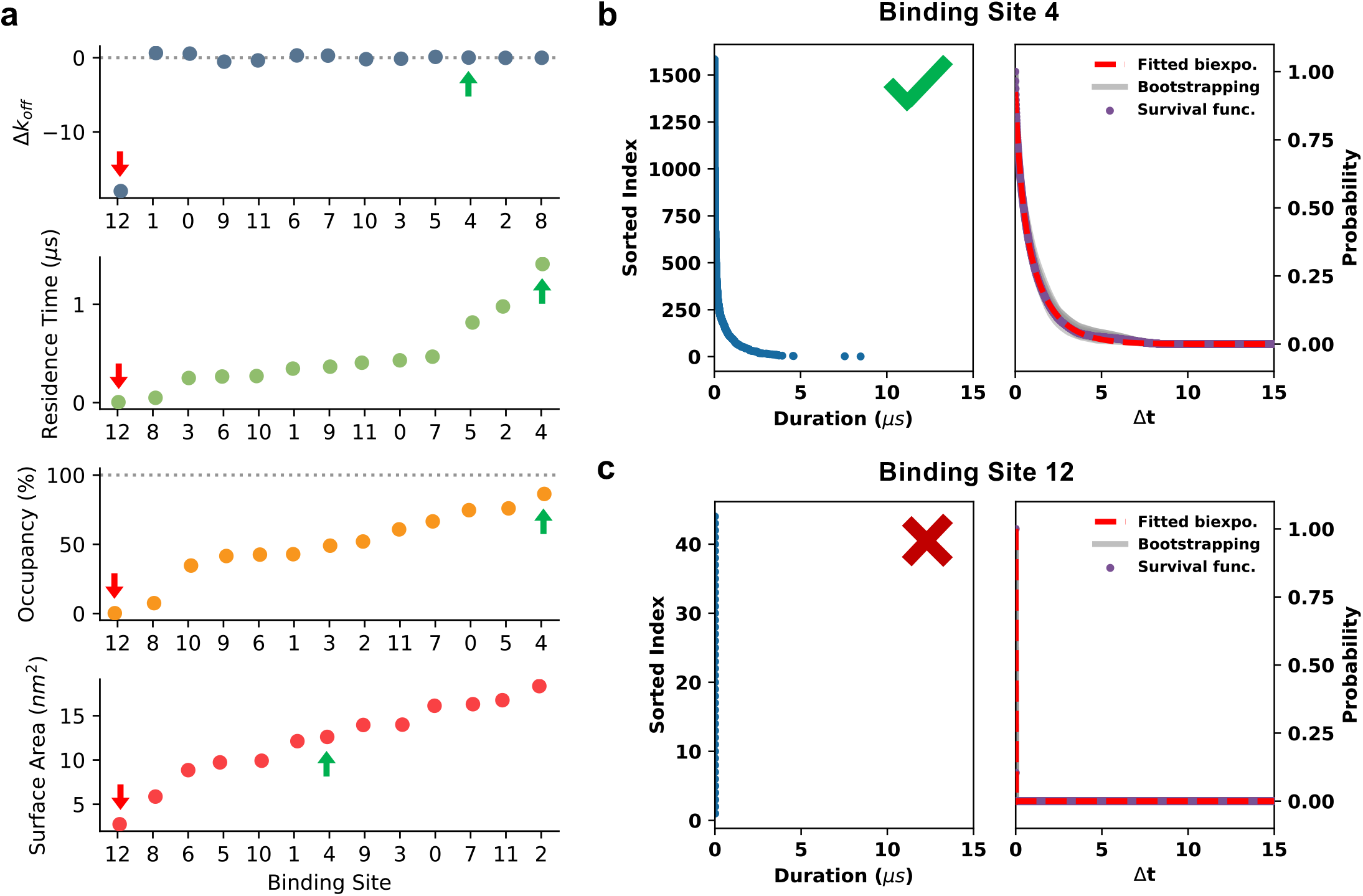
Screening binding site data. Metrics for discerning binding site quality during processing of PyLipID outputs. **a)** Comparison of binding site Δ*k_off_* values (*k_off_* bootstrap – *k_off_* curve fit), residence times, site occupancies and surface areas for PIP_2_ interactions with HHAT (10 x 15 μs CG simulations). Binding sites are ranked either from lowest to highest (residence times/occupancies/surface areas) or from worst agreement between calculated site *k_off_* values (Δ*k_off_*) to best (i.e., closest to 0). Arrows indicate sites corresponding to those in **b** (green) and **c** (red). **b-c)** Example binding site plots for PIP_2_ binding to a **b)** well sampled site (BS4) and **c)** an infrequently observed site (BS12) on HHAT. In each case a sorted index of interaction durations within the simulations is shown on the left panel. The right plot corresponds to the survival time correlation function of interaction durations (blue dots). *k_off_* values are derived either via biexponential curve fitting to the survival time correlation function (red line) or via bootstrapping (grey lines).

### Comparing lipid poses with cryo-EM densities

Subsequent stages of the pipeline concern simulation-assisted interpretation of structural lipid-like densities. For HHAT, we compared the top ranked CG lipid binding poses with the position of cryo-EM densities and ranked the relative residence times of all lipids binding to the same site (Fig. 4). These plots can be used to assess how binding site properties may dictate binding of a particular lipid type and evaluate the relative specificity of the site. For example, a site of lipid tail-insertion within HHAT (Fig. 4a) shows equivalent preference for PC and PE lipids whereas a surface site (Fig. 4d) preferentially binds anionic lipids. Refinement of lipid binding poses using atomistic simulations revealed remarkably good overlap with densities, quantified by Q scores^25^ for the lipid poses (Qavg = ~0.4 compared to ~0.7 for structurally modelled palmitate moieties and HHAT heavy atoms at 2.7 Å) (Fig. 4, Extended Data Fig. 2). This is particularly impressive considering lipid poses were derived *ab initio* from the simulations and in the absence of any density guided restraints. We note that LipIDens can be employed iteratively throughout the model building process, including for low-resolution maps. We exemplify this for HHAT using a low-resolution map at ~5 Å (Fig. 4a) whereby PyLipID was able to identify a lipid binding site corresponding to kinked tail density which was subsequently revealed (among the other peripheral densities) when the map resolution was improved to ~2.7 Å (Fig. 4b-e), thus serving as a doubleblind test study.

**Figure 4:**
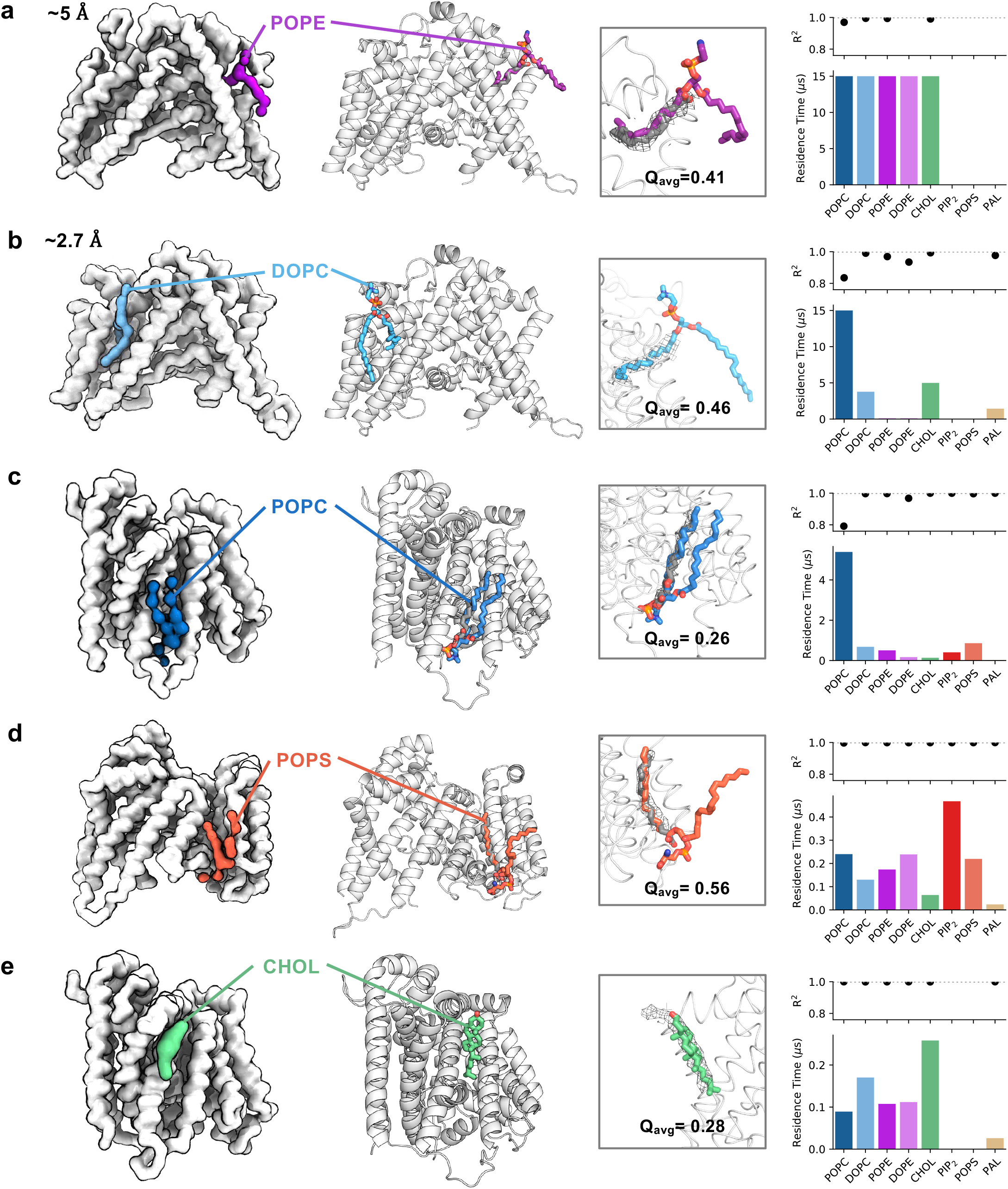
Comparison of cryo-EM densities with lipid poses from simulations. Identification of representative bound poses of lipid species to assist interpretation of cryo-EM densities, exemplified for lipid interactions surrounding HHAT. Left: CG binding poses for lipids bound to identified binding sites on HHAT. CG simulations were initiated using a low-resolution structure derived from a preliminary cryo-EM map (**a**, ~5 Å) or a higher resolution map (**b-e**, ~2.7 Å)^27^ to illustrate how LipIDens can be implemented throughout the model building process. HHAT was simulated for 10 x 15 μs in each case. Middle left: selected pose of a lipid bound to HHAT during atomistic simulations initiated by back-mapping from CG simulations. Middle right: comparison of cryo-EM densities (grey mesh) with the atomistic pose. Modelled palmitate moieties in the HHAT structure are shown as grey sticks. Average Q scores^25^ for the atomistic lipid tail pose within the cryo-EM density are indicated. Right: binding site residence times and R^2^ values for each lipid which binds to the site, used to assess preferential binding of a lipid species to specific sites. POPC is coloured dark blue, DOPC light blue, POPE purple, DOPE pink, cholesterol green, PIP_2_ red, POPS coral and palmitate (PAL) ochre throughout.

### Application to other membrane proteins

We applied the pipeline to three different membrane proteins for which lipids have been assigned to putative densities in recent structures; the eukaryotic proton channel Otopetrin1 (OTOP1)^34^, the *Escherichia coli* pentameric ligand-gated ion channel ELIC^35^ and the mechanosensitive channel of small conductance (MscS), also from *E. coli*^36^ (Fig. 5). These examples serve to demonstrate the diverse applicability of LipIDens to assist interpretation of structure-function questions.

**Figure 5:**
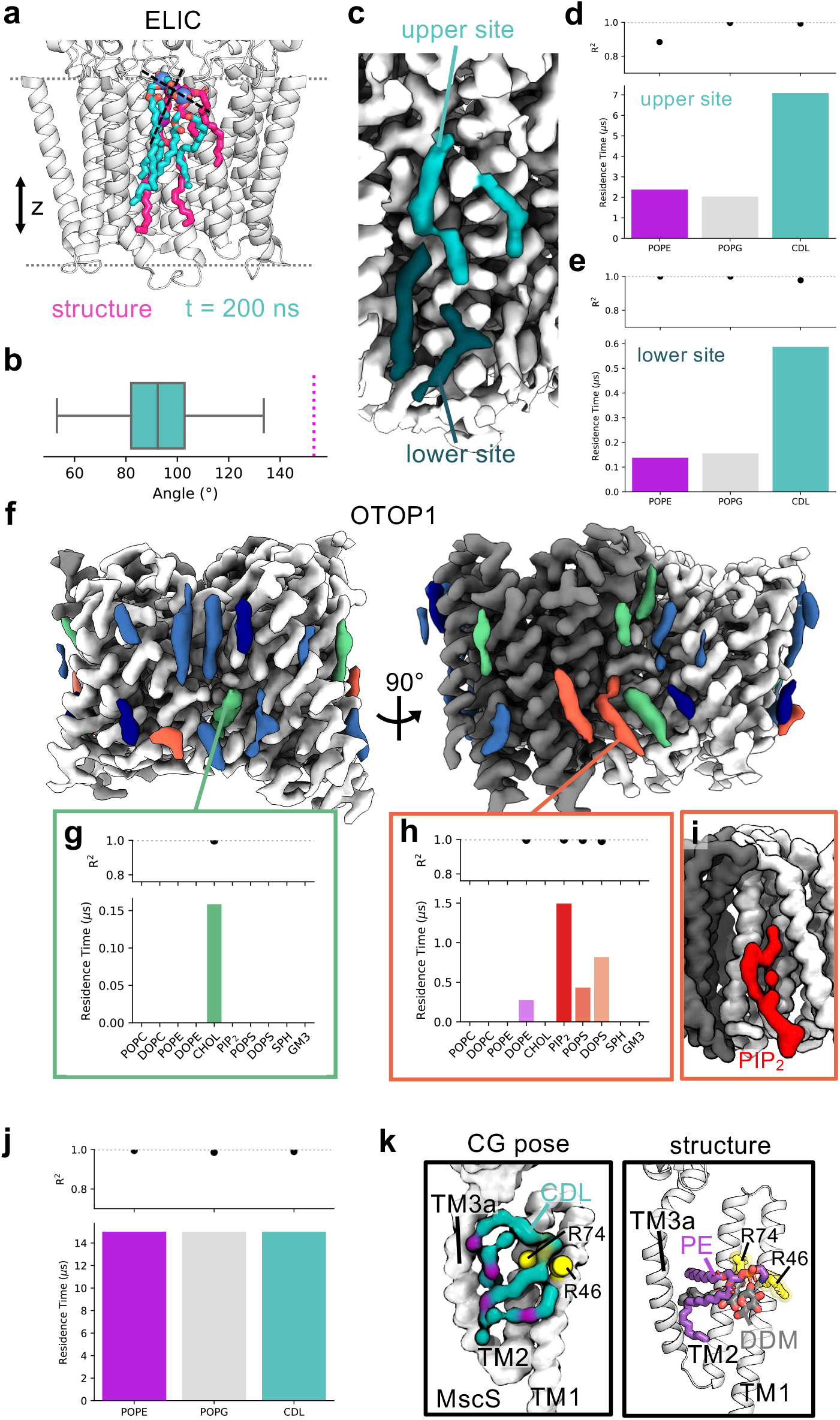
Application of the pipeline to a range of example proteins. The LipIDens pipeline as applied to assist interpretation of lipid-like densities within structures of **a-e**) the *E. coli* pentameric ligand-gated ion channel (ELIC, PDBid 7L6Q)^35^, **f-i**) the proton channel Otopetrin1 (OTOP1, PDBid 6NF4)^34^ and **j-k**) the *E. coli* mechanosensitive ion channel (MscS, PDBid 7ONJ)^36^. Each protein was simulated for 10 x 15 μs in a lipid composition designed to mimic the native environment of the protein (Supplementary Table 2). **a)** Overlay of the structurally modelled cardiolipin (CDL) pose on ELIC (magenta) with the pose at the end (t = 200 ns) of an atomistic simulation (teal) initiated from the top ranked CG CDL binding pose identified by PyLipID^24^. Phosphate groups of each CDL molecule are shown as spheres connected by a vector indicating the relative lipid tilt angle with respect to z. **b)** Angle of the vector between CDL phosphate groups with respect to z across 3 x 200 ns atomistic simulations (teal). The magenta line indicates the structurally modelled lipid tilt angle. **c)** Lipid-like densities at the proposed site indicating discontinuous density at the bilayer midsection to form two distinct CDL binding sites in the upper (teal) and lower (dark teal) leaflet. Site residence times and R^2^ values for PE, PG and CDL binding to the identified upper (**d**) and lower (**e**) sites. **f)** Lipid-like densities surrounding OTOP1 coloured according to whether cholesterol (green) or PIP_2_/PS (red) were identified as binding to the site among the highest lipid residence times. Other lipid densities where sites were identified by PyLipID are shown in blue (see Extended Data Fig. 4) and densities where lipid sites were not identified by PyLipID are shown in dark blue. **g)** Exclusive binding of cholesterol between the N- and C-domains of OTOP1, corresponding to the cholesterol site modelled in the structure^34^. **h)** Preferential binding of anionic lipids in proximity to a kinked lipid density at the OTOP1 dimer interface. **i)** Top ranked PIP_2_ binding pose identified by PyLipID from CG simulations, showing curved tail position which matches the lipid density at this site. **j)** Prolonged interactions of PE, PG and CDL at the lipid site on MscS between TM2 and TM3a. **k)** Comparison of the top ranked CDL binding pose from CG simulations (left) with the modelled PE and DDM molecules in the MscS structure (right) showing tail insertion/stacking between TM2 and TM3a and a tilted lipid binding pose.

In the ELlC structure, authors observe an elongated density traversing both leaflets, modelled as a highly unusual extended and tilted cardiolipin (CDL) molecule (Fig. 5a, magenta)^35^. In simulations we also observe CDL binding to this site, constituting the top ranked CDL site across the protein (Extended Data Fig. 3). We were unable to replicate the unusual tilted modelled pose despite pose refinement with atomistic simulations (Fig. 5a, teal). We observe a more conventional CDL binding pose whereby the phosphate beads remain in close z axial proximity (Fig. 5b, Extended Data Fig. 3), consistent with a large-scale analysis of CDL binding poses in *E. coli*^31^. Re-assessment of the proposed CDL density shows discontinuity at approximately the position of the bilayer midplane (Fig. 5c). Consistent with this we identified a second lipid site in the inner leaflet which also preferentially bound CDL, albeit with a much lower residence time (Fig. 5d-e). This raises the possibility that the density in fact corresponds to two lipids in adjacent leaflets, for which additional experimental analysis will be required to establish (Fig. 5d-e). The diffuse nature of densities in this region may also be accounted for by tail promiscuity/dynamics across the two CDL binding sites, a feature we also observed in atomistic simulations (Extended Data Fig. 3c). This highlights the highly non-trivial nature of interpreting lipid-like densities from cryo-EM structures and the power of the pipeline to assist model building and density interpretation.

For OTOP1, assignment of the putative lipid densities was challenging, due to resolution ranging 3.1-3.4 Å around the TMD. The authors assigned three densities per protein subunit as cholesterol-hemisuccinate (CHS), trapped between the dimer interface and thus occluded from the bilayer accessible region. An additional density between the N- and C-domain of each monomer was modelled as cholesterol^34^. Assignment of these densities was likely possible due to enclosure between the transmembrane segments which may have stabilised the bound lipids/detergents. Given these observations we used LipIDens to assess which of the remaining 17 densities per monomer may also correspond to cholesterol. Cholesterol binding poses matched the location of 4/17 of the additional lipid-like densities (Fig. 5f, green), for which cholesterol was one of the highest ranked lipids (Extended Data Fig. 4). We were able to recapitulate exclusive binding of cholesterol at the N/C domain interface, consistent with the modelling in the structure (Fig. 5g). Modelling of this density as cholesterol is also ranked highly in the PDB ligand validation tool. In addition, we were able to use the pipeline to suggest the mostly likely identity of lipid species at those sites where cholesterol did not bind (Extended Data Fig. 4). We observed preferential binding of lipids with anionic headgroups (PIP_2_/PS) to three of these sites (Fig. 5f, red, Extended Data Fig. 4). This included one notable curved tail-like density at the edge of the dimer interface which was also captured in the top ranked PIP_2_ pose at this site (Fig. 5h-i). These densities may therefore correspond to bound PIP_2_ and/or PS molecules extracted from the native bilayer. There were 3 densities per monomer which we could not assign to lipids based on the top ranked simulation poses (Fig. 5f, dark blue, Extended Data Fig. 4). These densities were smaller and may result from differences between the binding properties of detergents *vs*. lipids or from the limited resolution of low occupancy binding events.

A high-resolution structure of MscS was solved to 2.3 Å allowing for modelling of 8 detergent moieties per subunit (5x lauryl maltose neopentyl glycol (LMNG), 3x N-dodecyl-β-maltoside (DDM)). The authors were also able to resolve a bound lipid, assigned as PE, which was tilted by ~80° degrees with respect to the bilayer normal^36^. We wished to assess whether a) PE preferentially bound to this site when MscS was embedded in an *E. coli* inner membrane-like lipid composition (i.e. PE/PG/CDL) and b) whether a tilted lipid conformation was also observed when the protein is embedded within a lipid bilayer. In simulations, this site emerges as a prominent and prolonged binding site for PE, PG and CDL with all lipid types binding with residence times of at least 15 μs (Fig. 5j). This is consistent with an experimental study suggesting the pocket can be accessed by multiple lipid types, including CDL, in a manner that was broadly independent of the headgroup type^37^. Assessment of the top ranked lipid binding poses revealed a tilted conformation for CDL with the tails inserting into a groove between TM2 and TM3a and the phosphate headgroups coordinated by R46 and R74 (Fig. 5k, Extended Data Fig. 5). This also highlights the ability of simulations to provide additional native context, given CDL was not added during determination of the MscS structure. We did not observe lipid tilt amongst the top ranked poses of PE or PG but tilted conformations were present in subsidiary pose clusters. The trapped CDL tail between TM2 and TM3a is intriguing since the acyl-tail of DDM is observed to occupy the same groove as the PE tails in the MscS structure (Fig. 5k, Extended Data Fig. 5). Thus, DDM may aid stabilisation of the protein by mimicking the behaviour of ‘bulkier’ lipid types with additional tails (such as CDL) in a detergent context and/or by displacing tail binding from the groove during protein solubilisation. It is also possible that DDM may modify the hydrophobic volume of the groove between TM2/TM3a to accommodate the tilted PE molecule.

## Discussion

In summary, we have developed the LipIDens pipeline for simulation-assisted interpretation and refinement of lipid-like structural densities. We describe how LipIDens can be applied to establish and analyse simulations and to assess the quality of lipid interaction data (Fig. 1-3). We detail how the pipeline can be employed to assess lipid site identity and specificity using HHAT as an example (Fig. 4). Finally, we assess lipid-like densities across a range of other membrane proteins to illustrate how LipIDens can be applied to:

1. Identify and refine lipid binding poses using a multiscale simulation approach (Fig. 5a-e).
2. Suggest the most likely identity of lipid densities and rank the relative residence times of different lipids binding at a site (Fig. 5d,e,g,h,j).
3. Differentiate between lipid-tail and sterol like densities (Fig. 5f).
4. Identify differences between structural densities and simulation derived lipid poses (Fig. 5f).
5. Discriminate between binary lipid binding sites and those able to interact with a range of lipid types (Fig. 5f-i).
6. Capture possible occurrences of detergent biomimicry as exemplified by comparison of CDL poses with detergent/lipid stacking (Fig. 5k).

Cellular membranes contain hundreds of different lipid species, with highly diverse headgroup and tail compositions dependant on e.g. subcellular localisation^38–40^. Only a subset of these lipid types are available for use in CG simulations, although topology files for the most abundant lipid species are generally available^41^. Consequently, the goal of this pipeline is not to definitively identify exact molecular identity *per se* of a bound lipid at a site but to guide the user towards the most likely identity of the lipid within a given membrane composition. As such, selected membrane compositions should mimic, at least to a first approximation, the native environment of the membrane protein or experimental lipid conditions (such as the nanodisc composition)^42–45^. In particular, if there is already data suggesting a biological role for a specific lipid, it would of course be wise to include this in the bilayer component of the simulation. In addition we note there is likely to be some bias in the initial density map towards lipids with strong interactions which are able to survive membrane protein purification, as has been suggested by previous affinity calculations^46^.

One key feature of LipIDens is the ability to capture lipid binding sites and representative poses *a priori* from unbiased (equilibrium) simulations whereby, unlike in e.g. docking studies (where search space is restricted) sites are explored over the whole membrane lipid accessible surface. LipIDens also automates processing and validation steps to readily obtain meaningful results from these comprehensive data sets. Ultimately, the LipIDens pipeline demonstrates how integrative structural biology methods can be applied to facilitate the biologically relevant contextualisation of membrane protein structures.

## Methods

### Input data

Protein coordinate files in .pdb format and corresponding cryo-EM density map for the protein (e,g. from the Electron Microscopy Data Bank (EMDB) https://www.ebi.ac.uk/emdb/) are required. MARTINI (version 2.2 or 3.0) parameters (http://cgmartini.nl/index.php/downloads) are used for CG simulations and automatically obtained by LipIDens. For atomistic simulations, CG2AT provides a choice of forcefields automatically^48^. Molecular dynamics simulation parameter files are automatically provided in the pipeline. The default linear constraint solver (LINCS)^49^ parameters (lincs_order=4, lincs_iter=1) are used in GROMACS mdp files unless MARTINI-2.2 cholesterol with virtual sites^50^ is included in the bilayer, in which case lincs_order=12 and lincs_iter=2 are used instead, in line with recent findings^51^.

Molecular dynamics simulations in the examples described used GROMACS 2019 (> version 5 recommended) (https://www.gromacs.org/), with visualisation using VMD^52^ (https://www.ks.uiuc.edu/Research/vmd/) and PyMOL (https://PyMOL.org/2/). The LipIDens pipeline was installed from the GitHub repository (https://github.com/TBGAnsell/LipIDens). LipIDens uses additional packages which are automatically installed: PyLipID (version >=1.5)^24^ (from https://github.com/wlsong/PyLipID) and Martinize2 (version >=0.7) (https://github.com/marrink-lab/vermouth-martinize). Additionally, dssp (https://swift.cmbi.umcn.nl/gv/dssp/); CG2AT (https://github.com/owenvickery/cg2at)^48^; and propKa (https://github.com/jensengroup/propka)^53^ may be required.

### LipIDens Pipeline

The LipIDens pipeline is composed of multiple stages, run using an interactive standalone master python file (‘lipidens_master_run.py’) or by pre-defining variables, as described in the jupyter (https://jupyter.org) notebook (‘LipIDens.ipynb’). A detailed step-by-step guide to LipIDens usage is provided in the accompanying protocol XXX (https://protocolexchange.researchsquare.com). The GROMACS 2019 MD simulation software^54^ (https://www.gromacs.org/) was employed throughout. Additionally, the MARTINI-2.2 forcefield was used for CG simulations^41^ due to its broad applicability and ability to replicate experimentally observed lipid binding poses^55^. The protocol can also be used with MARTINI-3.0 if required.

#### Coarse-grained MD simulations

Simulations of HHAT were initiated using coordinates derived from two cryo-EM maps at ~2.7 Å (PDBid: 7Q1U)^27^ and ~5 Å resolution (unpublished). HHAT CG simulations were set up as described in ^27^ and as detailed in the accompanying protocol for all proteins. Coordinates for OTOP1 and ELIC were derived from the Protein Data Bank (PDB) (PDBid: OTOP1 6NF4, ELIC 7L6Q)^34,35^. The structure of MscS was kindly provided by Dr. Tim Rasmussen, and is now also obtainable from the PDB (PDBid: 7ONJ)^36^.

Simulations were setup as described in detail in the accompanying protocol (https://protocolexchange.researchsquare.com). The MARTINI-2.2 forcefield^41^ was used to describe all components and simulations were performed using GROMACS 2019^54^ (www.gromacs.org). Lipid compositions were selected to recapitulate the native bilayer composition of each protein (as detailed in Supplementary Table 2). Alternatively, LipIDens provides a number of default membrane compositions (Supplementary Table 1). Energy minimisation, equilibration and production simulations were run using the parameters detailed in the .mdp files within the GitHub repository. Each system was simulated for a total of 10 x 15 μs.

#### Testing PyLipID cut-offs

PyLipID analysis was used to test lower and upper cut-off values to define interactions of a specific lipid with a protein. In general, it is recommended to exhaustively test a range of upper and lower cut-off value pairs over a few different lipid types, particularly those which are chemically diverse such as e.g. sterols *vs*. phospholipids. The output from this analysis is provided as a plot of interaction duration times, number of calculated binding sites and number of contacting residues for each dual cut-off combination (Extended Data Fig. 1e-g). In addition, a probability distribution plot of minimum lipid-residue distances is also generated by LipIDens (Fig. 2a, Extended Data Fig. 1a-d).

Appropriate lower and upper cut-offs correspond approximately to the position of the first solvation peak and the proceeding trough respectively (Fig. 2a). In addition, the lower cut-off demarks the point at which there is a jump in calculated duration times, binding site numbers and contacting residues when exhaustively testing cut-off pairs. Choice of upper cut-off also depends on whether deviations are observed in the exhaustive cut-off search when the upper cut-off is changed. Ideally the interaction metrics should plateau when an appropriate upper cut-off value is reached (Extended Data Fig. 1e-g).

#### Selecting PyLipID input parameters and running PyLipID analysis

The next step of the LipIDens pipeline relates to the computation of lipid binding sites and associated interaction kinetics using PyLipID. The lipid atoms included in site calculations can be tuned based on the putative lipid densities present in the corresponding cryo-EM maps by for example, restricting to lipid headgroup atoms (Fig. 2b). The sites calculated here included all lipid atoms and implemented a 0.475/0.7 nm dual cut-off scheme for all proteins. In the case of protein oligomers, OTOP1 (dimer), ELIC (pentamer) and MscS (heptamer), lipid interactions were averaged over protein sub-units. All other PyLipID input parameters were kept at default settings (binding_site_size=4, n_top_poses=3 and n_clusters=auto). PyLipID outputs were automatically mapped onto protein structures provided in the input .pdb file. Top ranked lipid poses, pose clusters, per residue and site kinetics and structural coordinates with kinetics mapped to the B-factor column were generated by PyLipID.

#### Screening PyLipID data

LipIDens ranks the lipid binding sites generated by PyLipID from lowest to highest (in the case of e.g. Occupancy, Residence time or Surface area) or closest to 0 (for Δ*k_off_* where Δ*k_off_* is the difference between the *k_off_* calculated form the curve fit of the survival function and the bootstrapped *k_off_* of the same data) (Fig. 3a). Poorly defined sites with large Δ*k_off_* values (generally > ± 1 μs) were excluded from future stages of the pipeline. Site ranking was used to identify sites with long residence times and occupancies and with Δ*k_off_* ~ 0 μs which may be of biological relevance and/or for comparison with cryo-EM densities. It is useful to inspect the mean survival time correlation function plots to assess site sampling and quality of calculated binding sites (Fig. 3b-c). The interaction durations plots should be well populated and the biexponential fit/bootstrapping curves should approximate the underlying survival function data (Fig. 3b). Additional R^2^ values for predicted residence times are provided as a further metric for assessing the quality of PyLipID outputs. If most of the sites are not well defined, this is usually an indication you should increase the length of simulations to improve site sampling.

#### Comparing lipid poses with cryo-EM densities

Bound lipid poses outputted by PyLipID were visualized using VMD, for both the top ranked lipid binding poses (‘*BSidX_rank*’) and the clustered poses (‘*BSidX_clusters*’). The identified binding poses (excluding poses from poorly defined sites as identified previously) were compared with the position of densities in the cryo-EM maps.

#### Ranking lipid species at a site

LipIDens generates plots to compare the residence times and R^2^ values of different lipids binding to the same site (Fig. 4, Extended Data Fig. 4). LipIDens automatically calculates the closest matching binding sites for selected lipids based on similarity between binding sites residues. Residues comprising binding sites are compared to those of the reference lipid (i.e. the first lipid inputted when prompted). It is recommended to use an abundant phospholipid (rather than e.g. a sterol) as the reference lipid. These were further inspected manually to check predicted site matches and remove poorly defined sites. By comparing the binding poses of the reference lipid to the location of lipid-like densities in the cryo-EM map the plots can be used to infer the most likely identity of the lipid species accounting for a given density.

#### Lipid pose refinement using atomistic simulations

The final stage of the LipIDens pipeline generates inputs for atomistic simulations which can be used to refine the CG lipid poses. CG simulations frames (i.e. those from which the top ranked CG lipid poses were derived) were back-mapped to atomistic resolution using CG2AT^48^ which generates all inputs and parameters needed for simulation with GROMACS. Atomistic simulations of HHAT were performed as described for the apo state (5 x 200 ns) in ^27^ and detailed within the accompanying protocol. Additional atomistic simulations (8 x 200 ns) were established via back-mapping from different CG frames to refine the poses of different lipids. Setup of the additional simulations was performed identically to previous replicates. For ELIC the CG frame from which the top ranked cardiolipin binding pose was derived was backmapped to atomistic resolution, energy minimised and equilibrated using CG2AT^48^. The CHARMM-36 forcefield^56^ was used describe all components and simulations were performed using GROMACS 2019^54^ (www.gromacs.org). The ELIC system was simulated for 3 x 200 ns. Parameters used in the production run are provided in .mdp files on the GitHub page (CG: https://github.com/TBGAnsell/LipIDens/tree/main/lipidens/simulation/mdp_files, atomistic: https://github.com/TBGAnsell/LipIDens/tree/main/lipidens/simulation/mdp_files_AT).

Once the atomistic simulations had finished running, refined lipid binding poses were compared to the cryo-EM density. The match between a simulation derived lipid pose and the cryo-EM density can be evaluated using Q scores^25^ within in UCSF Chimera using the MapQ plugin^25^. Average Q scores of lipid tails were calculated for HHAT in regions overlaying the density (Fig. 4), along with corresponding per atoms values (Extended Data Fig. 2). We note that low Q score values are calculated for lipid regions outside densities, consistent with increased lipid fluctuation of these exposed regions (Extended Data Fig. 2).

## Reporting Summary

### Data availability

Simulation parameter files compatible with GROMACS (*.mdp files) are embedded within the LipIDens pipeline and accessible on the GitHub page (CG: https://github.com/TBGAnsell/LipIDens/tree/main/lipidens/simulation/mdp_files, atomistic: https://github.com/TBGAnsell/LipIDens/tree/main/lipidens/simulation/mdp_files_AT).

Forcefield parameters compatible with MARTINI are automatically obtained by LipIDens from http://cgmartini.nl. Atomistic parameters are from CG2AT (https://github.com/owenvickery/cg2at).

The accompanying LipIDens protocol is provided at XXX (https://protocolexchange.researchsquare.com).

### Code availability

The LipIDens pipeline and codes described within this work are available at https://github.com/TBGAnsell/LipIDens. Notebook workflows (LipIDens.ipynb) and python scripts (lipidens_master_run.py) to run LipIDens are found on the GitHub page.

### Author contributions

T.B.A. implemented the LipIDens pipeline, ran and analysed simulations. W.S., R.A.C. and A.L.D were involved in method development for the PyLipID software. R.A.C, C.K.C. and M.M.G.G. tested the code. C.E.C., L.C., C.S., T.R. and A.B.W. provided structures for simulation and experimental expertise on cryo-EM processing. P.J.S. and M.S.P.S. conceptualised the project. T.B.A. wrote the paper and all authors provided comments and edited the manuscript.

## Acknowledgements

We thank Dr Zachary Berndsen for helpful discussion when preparing the manuscript. The LipIDens logo was designed by Jessica Ansell. T.B.A, C.E.C and M.M.G.G. are supported by Wellcome (102164/Z/13/Z). W.S. acknowledges support from the Newton International Fellowship. R.A.C., A.L.D. M.S.P.S. and P.J.S. are funded by Wellcome (208361/Z/17/Z). A.L.D. has been additionally supported by BBSRC (BB/R00126X/1) and the Department of Biochemistry. C.K.C. is funded by the BBSRC (BB/S003339/1). P.J.S.’s laboratory is also supported by the BBSRC (BB/P01948X/1, BB/R002517/1, and BB/S003339/1), and the MRC (MR/S009213/1) and M.S.P.S.’s research is further supported by the BBSRC (BB/R00126X/1) and PRACE (Partnership for Advanced Computing in Europe, 2016163984). C.S. is supported by Cancer Research UK (C20724/A26752), the BBSRC (BB/T01508X/1) and the European Research Council (647278). L.C. is supported by a Wellcome administrative support grant (203141/Z/16/Z). A.B.W. acknowledges support from the Ray Thomas Edwards Foundation. We thank Dr Irfan Alibay and Michael Horrell for the maintenance of local compute resources.

## Additional Information

Supplementary Information are provided.

## Extended Data Figure Legends

**Extended Data Figure 1:**
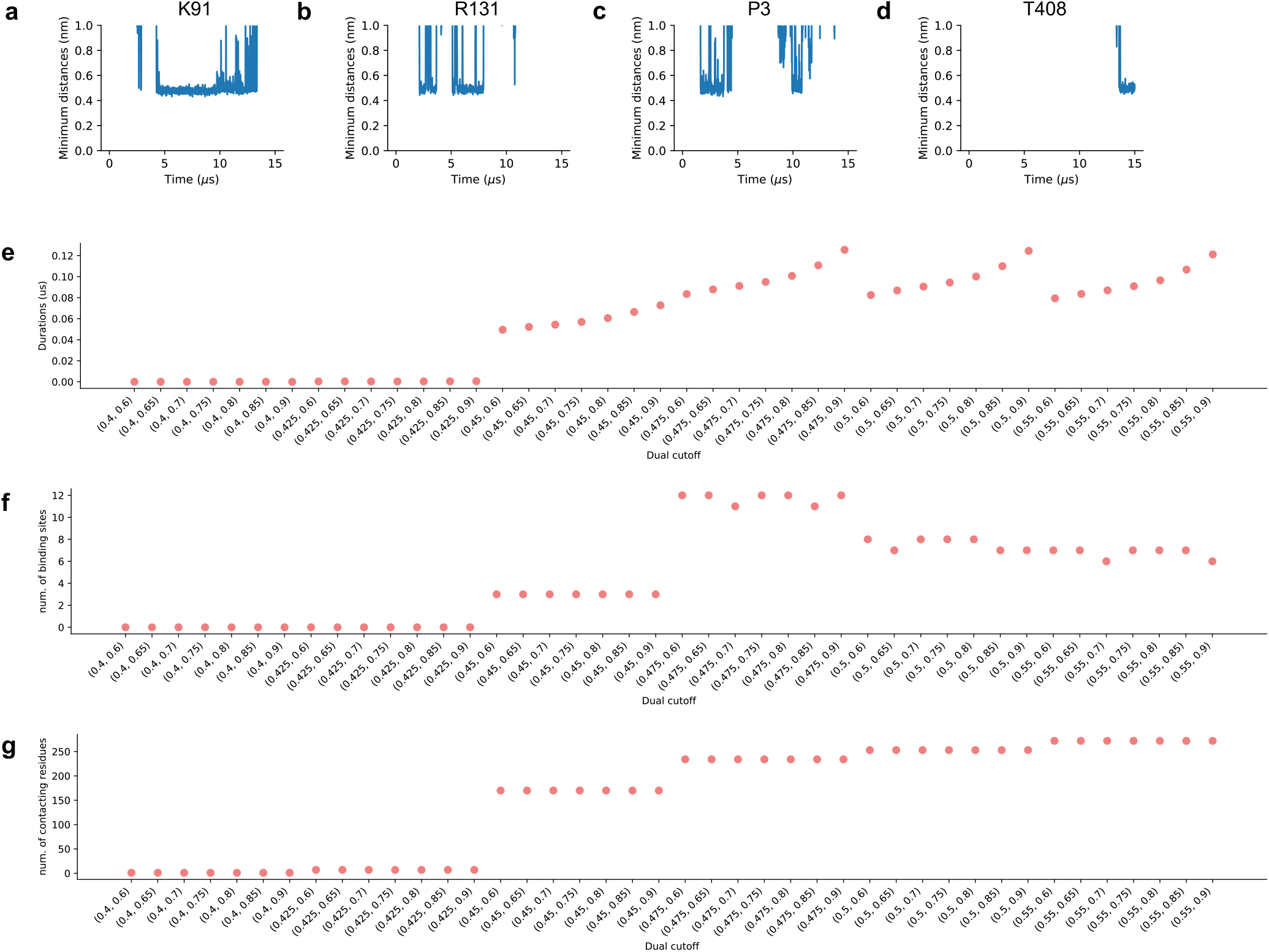
Tuning PyLipID cut-off values: interactions of HHAT with PIP_2_. Plotted outputs from PyLipID cut-off testing. **a-d**) Minimum distances between HHAT residues **a**) K91 **b**) R131 **c**) P3 and **d**) T408 and a PIP_2_ molecule across one 15 μs CG simulation. The minimum distance was calculated between any bead of the residue and any bead of the lipid. For clarity, only those interactions which came within 0.65 nm (distance_threshold) for at least 30 frames (contact_frames) of the simulation are plotted. **e-g**) Exhaustive testing of a range of lower and upper cut-off combinations for HHAT-PIP_2_ interactions (10 x 15 μs CG simulations). Plots show the effect of the selected cut-offs on **e**) interaction duration times **f**) the number of calculated binding sites and **g**) the number of interacting residues.

**Extended Data Figure 2:**
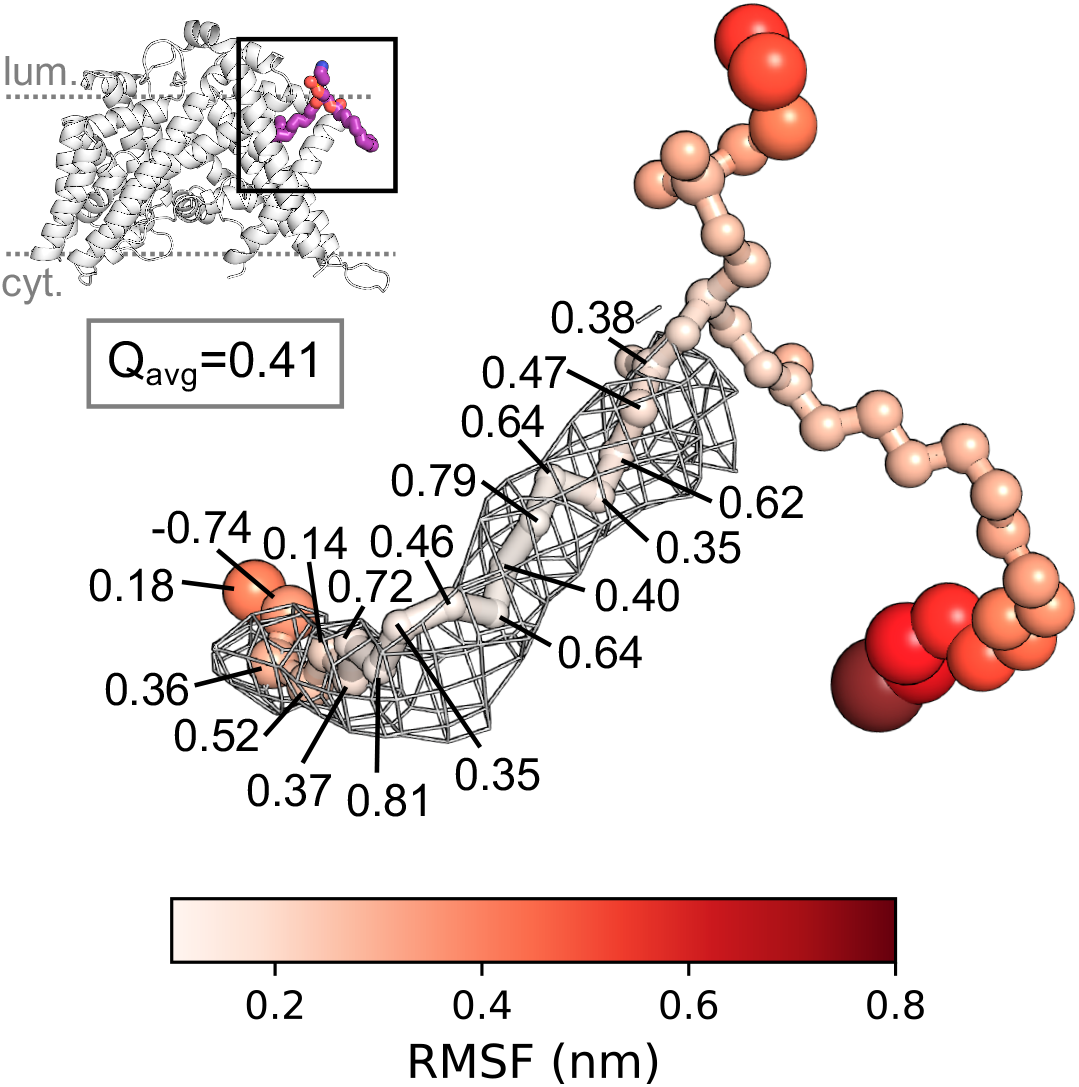
Comparison of lipid fluctuation with the cryo-EM density. The per atom root mean square fluctuation (RMSF) of a POPE lipid bound to HHAT (boxed) across 5 x 200 ns atomistic simulations. POPE atom spheres are scaled by RMSF value and coloured from low (white) to high (red). The per atom Q score^25^ was used to assess how well the simulation derived lipid pose matched the cryo-EM density.

**Extended Data Figure 3:**
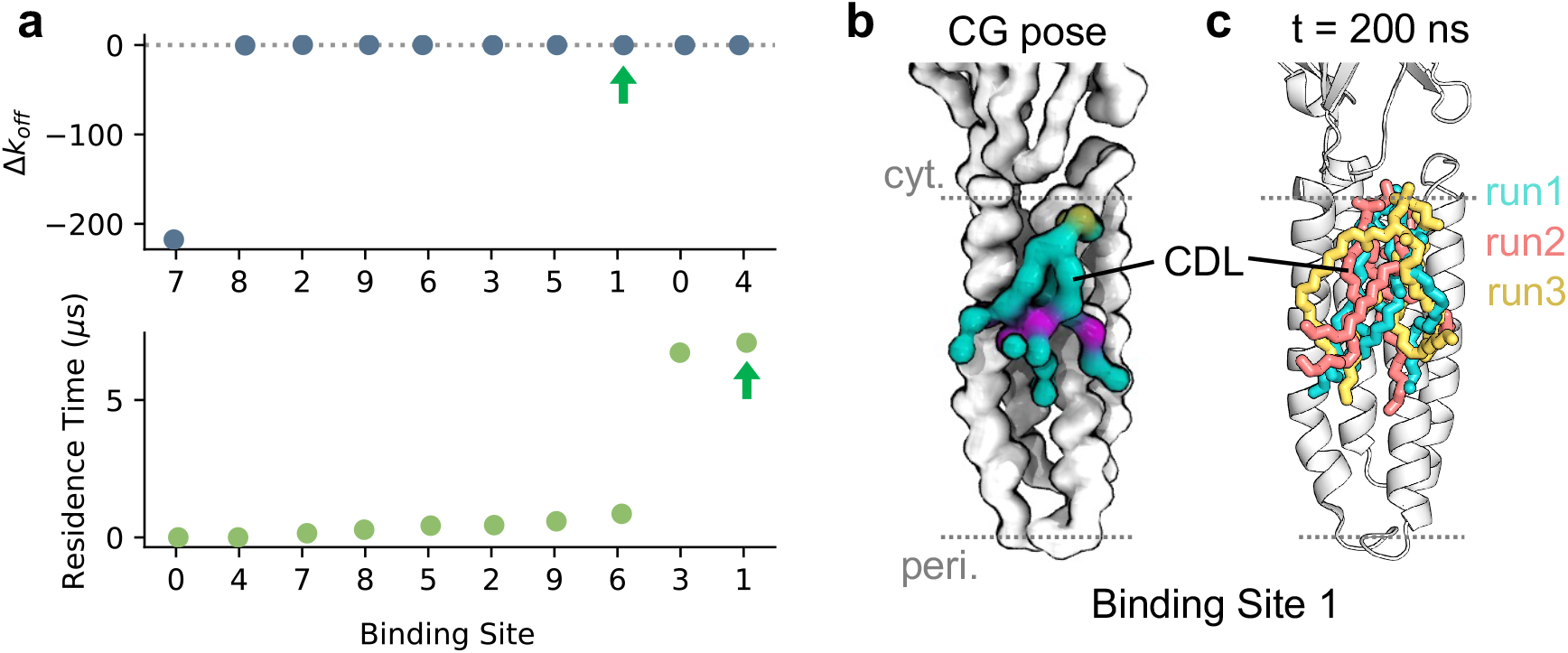
Cardiolipin binding to ELIC. **a)** Cardiolipin (CDL) binding sites ranked from worst to best Δk_off_ (Δk_off_ = k_off_ from curve fitting – bootstrapped k_off_) or lowest to highest residence time. The CDL binding site with the longest residence time, Binding Site 1, is arrowed. **b)** Top ranked CG binding pose for CDL at Binding Site 1. **c)** Snapshots of the CDL binding pose at the end of 3 x 200 ns atomistic simulations initiated using the CG CDL binding pose in **b**.

**Extended Data Figure 4:**
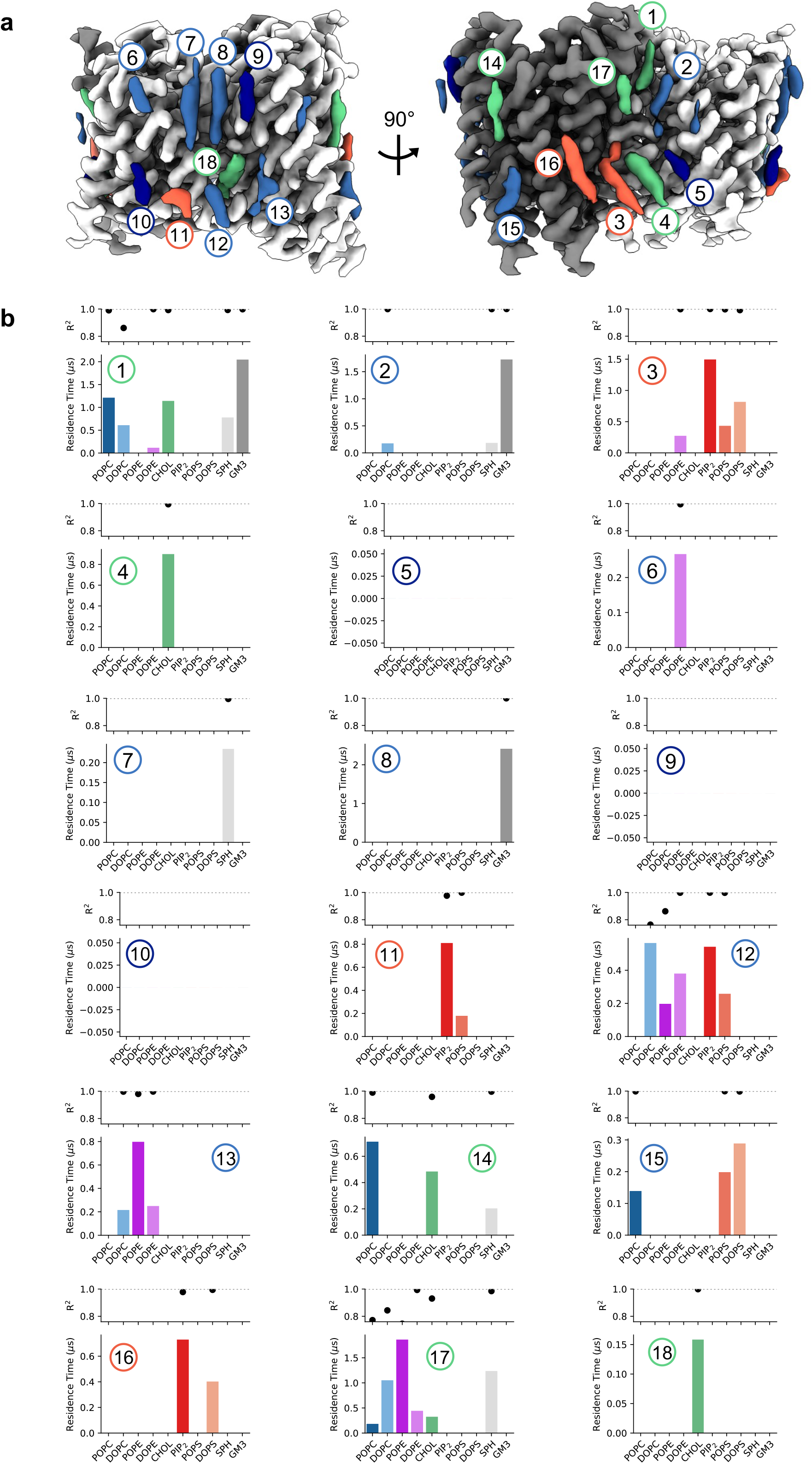
Interpretation of lipid densities surrounding OTOP1. **a)** Numbered lipid-like densities surrounding OTOP1. **b)** Residence time of lipids bound at sites corresponding to each numbered density. Lipid binding sites and residence times were calculated using PyLipID and a 0.475/0.7 nm cut-off from 10 x 15 μs CG simulations of OTOP1.

**Extended Data Figure 5:**
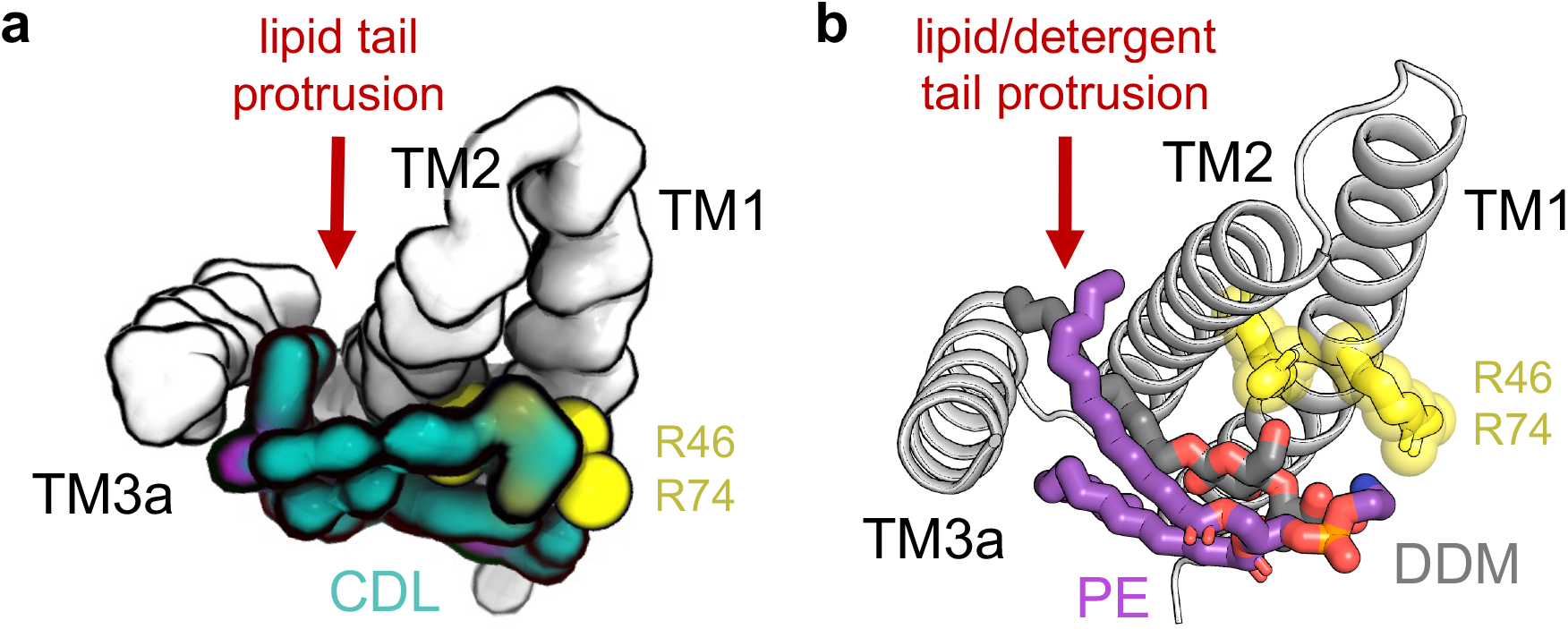
Lipid/detergent tail protrusion between TM helices of MscS. **a)** Cardiolipin (CDL) binding pose from CG simulations of MscS (viewed from the cytosolic membrane). The transmembrane helices of one monomer of MscS are shown as white surface and the position of residues coordinating the CDL headgroup are shown as yellow spheres. CDL tail protrusion between TM2/TM3a is arrowed. **b)** Structure of MscS (PDBid 7ONJ)^36^ in cartoon representation. The position of bound lipid (PE, purple) and detergent (DDM, grey) tails between TM2/TM3a is arrowed. Headgroup coordinating residues are shown as yellow sticks.

## Supplementary information

**Supplementary Table 1:** Predefined lipid compositions.

| Predefined bilayer type | CG bilayer composition (*insane.py* format)<sup>1</sup> |
| --- | --- |
| Simple | -u POPC:70 -u CHOL:30 -l POPC:70 -l CHOL:30 |
| Plasma membrane | -u POPC:20 -u DOPC:20 -u POPE:5 -u DOPE:5 -u DPSM:15 -u DPG3:10 -u CHOL:25 -l POPC:5 -l DOPC:5 -l POPE:20 -l DOPE:20 -l POPS:8 -l DOPS:7 -l POP2:10 -l CHOL:25 |
| ER membrane | -u POPC:37 -u DOPC:37 -u POPE:8 -u DOPE:8 -u CHOL:10 -l POPC:15 -l DOPC:15 -l POPE:20 -l DOPE:20 -l POPS:10 -l POP2:10 -l CHOL:10 |
| Raft-like microdomain | -u DPPC:27 -u DPPE:8 -u DPSM:15 -u DPG3:10 -u CHOL:40 -l DPPC:15 -l DPPE:35 -l DPPS:10 -l CHOL:40 |
| Gram neg. inner membrane | -u POPE:67 -u POPG:23 -u CDL2:10 -l POPE:67 -l POPG:23 -l CDL2:10 |
| Gram neg. outer membrane | -u PGIN:100 -l POPE:90 -l POPG:5 -l CDL2:5 |

**Supplementary Table 2:** Lipid compositions used in CG simulations.

| Protein | Native bilayer | CG bilayer composition |
| --- | --- | --- |
| HHAT | Endoplasmic reticulum | Upper: POPC (35%), DOPC (35%), POPE (8%), DOPE (7%), cholesterol (10%), palmitate (PCN) (5%)<br>Lower: POPC (15%), DOPC (15%), POPE (19%), DOPE (18%), POPS (8%), PIP<sub>2</sub> (10%), cholesterol (10%), palmitate (PCN) (5%) |
| OTOP1 | Plasma membrane | Upper: POPC (15%), DOPC (15%), POPE (2.5%), DOPE (2.5%), sphingomyelin (DPSM) (22%), GM3 (10%), cholesterol (33%)<br>Lower: POPC (7.5%), DOPC (7.5%), POPE (15%), DOPE (15%), POPS (7.5%), DOPS (7.5%), PIP<sub>2</sub> (7%), cholesterol (33%) |
| ELIC | *E. coli* inner membrane | POPE (67%), POPG (23%), cardiolipin (CDL2) (10%) |
| MscS | *E. coli* inner membrane | POPE (67%), POPG (23%), cardiolipin (CDL2) (10%) |

## Materials

### Expertise needed to implement the protocol

To implement the protocol a user should be familiar with basic terminal commands and navigation. Prior experience with setting up and running CG or atomistic MD simulations using e.g. GROMACS (www.gromacs.org) is advisable, for which tutorials are available elsewhere (http://www.mdtutorials.com/gmx/, http://cgmartini.nl/index.php/tutorials).

### Equipment

#### Input data

- Protein coordinate file in .pdb format as obtained either structurally or e.g. from the PDB (https://www.rcsb.org/). Protein models are also permitted and may be useful for switching to a different homologue (e.g. https://alphafold.ebi.ac.uk)^1^.
- Corresponding cryo-EM density map for the protein (locally determined or downloaded from the Electron Microscopy Data Bank (EMDB) (https://www.ebi.ac.uk/emdb/).

#### Optional input files

- Topology files for the system. Coarse-grained MARTINI parameters (http://cgmartini.nl/index.php/downloads) are automatically downloaded in the protocol or alternative parameters can be provided by the user. For atomistic simulations, CG2AT provides a choice of forcefields automatically^2^.
- Molecular dynamics parameter files. Simulation parameter files (e.g. mdp files for use with GROMACS) are automatically provided in the pipeline but users may wish to supply alternatives. The default linear constraint solver (LINCS) parameters (lincs_order=4, lincs_iter=1) are used in mdp files unless MARTINI-2.2 cholesterol with virtual sites^3^ is included in the bilayer, whereby lincs_order=12 and lincs_iter=2 are used instead, in line with recent findings^4^.

#### Computational hardware

- A computer or laptop with an operating system capable of running a terminal (Linux or MacOS recommended). For running simulations, a desktop computer, computational cluster or local high-performance computing facility is recommended for improved running times. Alternatively, national/cloud compute resources can also be used to run simulations.

#### Computational software

- Molecular dynamics simulation software for example GROMACS (> version 5) (https://www.gromacs.org/).
- Software for density map visualisation for example Coot^5^ (https://www2.mrc-lmb.cam.ac.uk/personal/pemsley/coot/), UCSF Chimera (https://www.cgl.ucsf.edu/chimera/), ChimeraX^6^ (https://www.rbvi.ucsf.edu/chimerax/) or PyMOL (https://PyMOL.org/2/).
- Software for trajectory visualisation (optional) e.g VMD^7^ (https://www.ks.uiuc.edu/Research/vmd/) or PyMOL (https://PyMOL.org/2/).
- python 3 (https://www.python.org/downloads/) (version >=3.9 recommended).
- The LipIDens pipeline (https://github.com/TBGAnsell/LipIDens). LipIDens can be installed by following the instructions of the GitHub repository. LipIDens uses the following additional packages which are **automatically** installed (see https://github.com/TBGAnsell/LipIDens/blob/main/requirements.txt for updated list of package dependencies):

○ PyLipID (version >=1.5)^8^. PyLipID is also available from https://github.com/wlsong/PyLipID and can be installed using pip install pylipid or git clone git://github.com/wlsong/PyLipID.git. PyLipID also implements the following additional packages (installed automatically): kneebow, logomaker, matplotlib>=3.3.4, mdtraj, networkx, numpy, p-tqdm, pandas, python-louvain, scikit-learn, scipy, seaborn, statsmodels, tqdm.
○ Martinize2 (version >=0.7) (https://github.com/marrink-lab/vermouth-martinize). Martinize2 can also be installed using pip install vermouth or pip install git+https://github.com/marrink-lab/vermouth-martinize.git#vermouth.
- dssp (https://swift.cmbi.umcn.nl/gv/dssp/).
- CG2AT (optional) (https://github.com/owenvickery/cg2at)^2^.
- propKa (optional) (https://github.com/jensengroup/propka)^9^.

## Procedure

LipIDens may be run either using an interactive standalone master python file (‘lipidens_master_run.py’) or by pre-defining variables within the pipeline jupyter (https://jupyter.org) notebook (‘LipIDens.ipynb’). The pipeline is composed of multiple stages, described under the subheadings below. When running LipIDens using the master python script, users are prompted to enter a number of variables. In the notebook, these are defined under the ‘USER DEFINED VARIABLES’ section, followed by the corresponding ‘CODE’ for each step.

Analysis stages of the protocol can be run independently if the user has pre-existing CG simulations they wish to examine, however we recommend some familiarity with all protocol steps.

The GROMACS 2019 MD simulation software^10^ (https://www.gromacs.org/) was used to run simulations. The MARTINI-2.2 forcefield was used for CG simulations^11^. The protocol could be adapted for use with other MD simulation software (e.g Amber, NAMD) or CG forcefields (e.g. MARTINI-3, ESPResSo^12^, Fat SIRAH^13^).

### Structure processing

TIMING: Steps 1-2, 5 mins, Step 3, 10-20 mins.

1. Load the input .pdb file in PyMOL and remove any superfluous components by selecting only the protein of interest and saving as a new .pdb file.
2. The input .pdb file should not have any residues with missing atoms but larger missing segments (e.g. loops, TM helices etc) are permitted. Simulation competent .pdb files of published membrane protein structures can be downloaded from MemProtMD (http://memprotmd.bioch.ox.ac.uk)^14^ or adjusted using https://github.com/pstansfeld/MemProtMD/blob/main/PDB-fix.ipynb. Decide whether any additional changes to the .pdb are required such as a) addition of missing atoms (required) or b) modelling of missing protein segments (optional) (step 3). If none, proceed to step 4.
3. Model missing atoms/segments using a preferred modelling software (MODELLER/Chimera-MODELLER^15^, SWISS-MODEL^16^, trRosetta^17^, AlphaFold2^1^, RoseTTAFold^18^).

### Setting up and performing coarse-grained simulations

TIMING: Step 4, 2 mins, Steps 5-6, ~2-10 days (variable depending on available compute resources, simulation system size and length)

4. Type python lipidens_master_run.py to initiate the LipIDens pipeline. The user will be prompted to select the directory to store simulations/data (save_dir) and the protocol stage (‘1a’). Default settings can be selected by pressing the ENTER/RETURN key. Follow the prompts to set a number of simulation variables described below. If using the jupyter notebook, define the variables for the first section of the protocol by modifying the file:

i. Change save_dir to a directory name in which to build the system and save analysis.
ii. Change protein_AT_full to the location of the input protein .pdb file.
iii. Set nprot to the number of homomeric protein chains or use nprot = 1 for heteromers.
iv. The variables protein_shift and protein_rotate can be used to alter the alignment of the protein within the bilayer. To alter the z axial position for protein insertion into the bilayer change the protein_shift value (decimal, negative and positive numbers accepted). Set the protein_rotate angle (in x y z) with respect to alignment along the first principal axis (default ‘0 90 0’ i.e. rotate in y by 90°). The position of the bilayer in MemProtMD (https://github.com/pstansfeld/MemProtMD/blob/main/MemProtMD_Self_Assembly.ipynb can be used to guide value selections^14^. ?TROUBLESHOOTING
v. Set the simulation boxsize (nanometres in x y z).
vi. Set the CG forcefield to use in simulations (currently compatibility with martini_v2.0, martini_v2.1, martini_v2.2 and martini_v3.0.0)^11,19^.
vii. The membrane_composition can be defined using either a) a predefined bilayer type or b) a custom bilayer composition. Current predefined bilayer names (e.g. ‘Plasma membrane’) and corresponding compositions are provided on the GitHub page. To build a custom bilayer change the membrane_composition variable to the chosen composition in the upper (-u) and lower (-l) leaflets using insane.py syntax^20^. For example, for a bilayer composed of 100% POPC in the upper leaflet and 90% POPC plus 10% POPS in the lower leaflet then membrane_composition=‘-u POPC:100 -l POPC:90 -l POPS:10’. Note, currently only a subset of lipids are available for the MARTINI-3 forcefield. Users should check whether the required lipids are available for a specific forcefield.
viii. Set martini_maxwarn to the maximum number of warmings permitted when running the martinize command (default 0).
ix. Change the ring_lipids variable to ‘True’ to place lipids within the protein when assembling the bilayer (default ‘False’).
x. Alter CG_simulation_time to the duration of each CG simulation in μs and set the number of replicates. It is recommended to run simulations for at least 10 μs and to run multiple repeats (at least 8 recommended) however the time taken to reach convergence will be system dependant and users should adjust accordingly. The ‘*Screening PyLipID data*’ section details how the quality of binding site data can be assessed using PyLipID outputs and, if necessary, users should consider extending the simulations.
xi. Finally alter n_cores to the number of CPUs to be used by GROMACS when setting up the CG simulations.
5. If using the notebook, run the code corresponding to CG simulation setup (up to the stage marked ‘PAUSE POINT’ after the ‘run_CG() function). This will happen automatically if using the master python script. The output of this step is a GROMACS md.tpr file for each CG simulation replicates. An outline of the commands **automatically** implemented within this section is given below:

i. Convert the protein to CG resolution using martinize2^21^ (https://github.com/marrink-lab/vermouth-martinize) with an ElNeDyn elastic network^22^ (for MARTINI-2) or Martini3001 forcefield (for MARTINI-3). A spring force constant of 1000 kJ mol^-1^ nm^-2^, a lower cutoff of 0.5 nm and an upper cut-off of 0.9 nm are applied. Users may alter the default elastic network, force constant and cut-off values within the lipidens/simulation/CG_simulation_setup.sh script using the -ff, -ef, -el, -eu, -ea and -ep flags. For more information see http://cgmartini.nl/index.php/tools2/proteins-and-bilayers/204-martinize.
ii. Embed the CG protein into a bilayer (of composition provided with the membrane_composition variable) and solvate with water using insane. py^20^. The position of the TMD within the bilayer is set using the protein_shift and protein_rotate variables. ?TROUBLESHOOTING.
iii. Neutralise the system by addition of ions using gmx grompp and gmx genion. By default, the system is neutralised using ~0.15 M NaCl however the cation and anion names and concentration can be altered using the -pname, -nname and -conc flags within CG_simulation_setup.sh.
iv. Make an index file (gmx make_ndx) which contains groups corresponding to the Protein, Lipids and Solvent (water and ions).
v. Generate a .tpr file for energy minimisation using gmx grompp and run the energy minimisation using gmx mdrun. ?TROUBLESHOOTING.
vi. Equilibrate the system using gmx grompp and gmx mdrun. In the first equilibration step restraints are applied to the protein backbone (BB) beads. In the second equilibration step restraint are not applied. ?TROUBLESHOOTING
vii. Generate the .tpr file for the production run using gmx grompp.
6. Run the CG simulation using gmx mdrun. ?TROUBLESHOOTING. It is recommended to offload this calculation to a cluster or high-performance computing facility. ·PAUSE POINT: wait until CG simulations have finished running before proceeding.

### Testing PyLipID cut-offs

TIMING: Step 7, 5-15 mins, Step 8, 2 mins, Step 9, ~1 h per lipid species, Step 10, 5 mins

7. Once all the CG simulations have reached completion, trajectories can be processed. Type python lipidens_master_run.py and select the protocol stage (‘1b’) when prompted, or run the corresponding notebook code. The trjconv_CG() function makes molecules whole across the periodic boundary and skips the number of frames provided with stride. Although not technically required for subsequent analysis, trajectory processing can improve the usability of outputted lipid binding poses from PyLipID and reduce PyLipID running times.

i. Set stride to the number of frames to skip during trajectory processing and downstream analysis of protein-lipid interactions (recommended to speed up processing).
8. Once the CG replicates are processed, PyLipID analysis can be performed. Δ CRITICAL STEP: In this first step, PyLipID is used to test a range of lower and upper cut-off values for lipid interactions with the protein. In general, users should test cut-offs for several chemically diverse lipids such as e.g. sterols or phospholipids. Run LipIDens and select the protocol stage (‘2’) when prompted. Within the notebook/master script set the user defined variables for the second section of the pipeline:

i. Set the lipid_atoms variable to the CG bead names PyLipID will use for cut-off testing. The default (lipid_atoms=None) will use all CG beads for each lipid.
ii. In the first stage of cut-off testing, the minimum distance of each lipid to a residue is plotted, provided that the lipid comes within distance_threshold of the residue for longer than the number of contact_frames. Set the distance_threshold value to a reasonably generous interaction distance (i.e. 0.65 nm for CG simulations or 0.4 nm for atomistic simulations). Select a value for contact_frames to screen interacting lipids.
iii. In the second stage of cut-off testing, a list of upper and lower cut-offs are exhaustively screened in a pairwise fashion. Change the lower_cutoff and upper_cutoff variables to lists of cut-off values to test (in nm).
iv. Change timeunit to the preferred axis unit on analysis plots.
9. Run the code corresponding to PyLipID cut-off screening (up until the next segment of user defined variables) within the notebook. This will happen automatically if using the python script. An outline of the steps implemented in this section are described below:

i. Calculate the minimum distances of each interacting lipid to a residue over the length of the trajectory. Plots are provided in the ‘*PyLipID_cutoff_test_Lipid/Figures*’ subdirectory.
ii. Plot the probability distribution of minimum distances between the lipid and the protein.
iii. Exhaustively test a range of upper and lower cut-off value pairs. The output is a plot of interaction duration times, number of calculated binding sites and number of contacting residues for each dual-cut-off combination.
10. Using the distribution plot from step 9ii and the exhaustive cut-off testing in step 9iii, select the lower and upper interaction cut-offs to use when running PyLipID.

i. The lower cut-off localises to the first solvation peak in the probability distribution plot. Additionally, the lower cut-off corresponds to an increase in interaction durations, computed binding sites and residues comprising each site compared with smaller lower cut-off values.
ii. The upper cut-off localises to the first trough between the first and second interaction shells in the probability density distribution. The upper cut-off is appropriate when interaction metrics plateau. If interaction metrics increase further as upper cot-off is increased this is an indication that the second solvation shell is being captured which should be avoided.

### Selecting PyLipID input parameters and running PyLipID analysis

TIMING: Step 11, 5 mins, Step 12, ~15 mins per lipid species

11. Next, lipid interactions, kinetics and binding sites are calculated using PyLipID. Run the LipIDens python script and select the appropriate stage (‘3’) when prompted. In the next user defined variables section of the notebook/master script set the following variables to run site analysis using PyLipID:

i. Set the cutoffs variable to the selected lower and upper cut-off (step 10). Additional information regarding cut-off selection is provided at https://pylipid.readthedocs.io/en/master/tutorials.
ii. Tune the lipid_atoms variable based on the putative lipid densities present in the cryo-EM map. If only headgroup-like density is present the lipid_atoms variable can be restricted to the CG headgroup beads to speed up calculation times. If tail density is present it is recommended to perform calculations on all lipid atoms however the search could be restricted to beads comprising the tail if required. This can be useful for assessing the relative contribution of different lipid segments to binding site residence times.
iii. If multiple, identical proteins and/or protein complexes are present, such as in homo-oligomeric ion channels, set the nprot flag to the copy number in the system. This will average calculated kinetic parameters over repeat domains and improve protein-lipid contact sampling.
iv. Set the binding_site_size valuable to the minimum number of residues that can comprise an identified binding site (default 4). This is recommended to avoid artefactual identification of very small ‘binding sites’ due to non-specific interactions.
v. Select the number of top lipid binding poses to be outputted for each binding site using the n_top_poses variable (default 3). At each site the specified number of representative lipid binding poses will be calculated using an empirical scoring function to rank lipid binding sites against the simulation derived lipid density at the site.
vi. Alter the n_clusters variable to calculate the number of distinct lipid pose clusters to be calculated for each site. This can be useful for assessing the conformational diversity of lipid binding poses at a site. If n_clusters is set to auto (default) PyLipID will use a density-based clustering algorithm to identify all possible clusters.
vii. Set save_pose_format to the coordinate file format for outputted lipid poses.
viii. Set save_pose_traj to True to output lipid binding poses to a trajectory format provided with save_pose_traj_format.
ix. Set the timeunit to use in outputted data.
x. Alter the resi_offset to offset the residue index number in outputs.
xi. The radii variable should be used to set the Van der Waals radius of non-standard atoms in a trajectory. The Van der Waals radius of common atoms are already accounted for, including CG beads in MARTINI 2.0-2.2.
xii. Set the pdb_file_to_map variable to an atomistic protein coordinate file (such as from step 4ii) onto which binding site information will be mapped by PyLipID.
xiii. Set the fig_format variable to the preferred image output file extension.
xiv. Change num_cpus to the number of CPUs to use during multiprocessing steps of PyLipID.
12. Perform PyLipID analysis on lipids in the CG simulations by running the corresponding section of code in the notebook. PyLipID runs automatically if using the master script. It is worth noting that PyLipID is highly modular and contains a number of functions that can be run independently to study other biological phenomena. Please refer to https://pylipid.readthedocs.io/en/master/ for details on how to write custom analysis scripts and/or select only those outputs of interest. PyLipID creates an ‘*Interaction_Lipid*’ directory containing the outputs for each lipid. This includes a ‘*Dataset_Lipid*’ directory containing data stored in pickle format and a summary of the kinetics associated with each residue and binding site (Dataset.csv). This subdirectory also includes a PyMOL script for automatically mapping binding site kinetics onto the atomistic structure provided with pdb_file_to_map. Other outputs include top ranked and/or clustered binding poses for each binding site (within the ‘*Bound_Poses_Lipid*’ subdirectory) and .pdb files with kinetics mapped to the B-factor column (within the ‘*Coordinate_Lipid*’ subdirectory).

### Screening PyLipID data

TIMING: Steps 13-16, 5 mins

13. The next stage of the pipeline involves inspecting and screening the PyLipID outputs. Within the notebook/master python file run the next section of code (‘4’) to rank binding site kinetics.
14. Inspect the output plot (Site_stats_rank_compare.pdf) located within the ‘*Interaction_Lipid*’ directory. The script ranks lipid binding sites from lowest to highest Occupancy, Residence time or Surface area or Δ*k_off_* closest to 0 (defined as the difference between the *k_off_* calculated form the curve fit of the survival function and the bootstrapped *k_off_* of the same data). This plot can be used to inspect the quality of calculated binding sites by e.g. comparing sites which rank highly (in their Residence times/Occupancies) and are well fitted/sampled. Typically, a good site has a Δ*k_off_* between ± 1 μs. ΔCRITICAL STEP: Always inspect the quality of the identified binding sites and remove any poor sites from future analysis.
15. Review the binding site *k_off_* plots (BS_idX.pdf) located within the ‘*Interaction_Lipid/Binding_Sites_koffs_Lipid*’ directory. Well sampled binding sites which rank highly should show good agreement between the bootstrapped and bi-exponential curve fits to the survival function and sufficient sampling of interaction durations. Poorly fitted sites are indicated by disagreement between bootstrapped and bi-exponential curve fits and/or sites which an infrequently observed as indicated by a sparse interaction duration plot. This serves as a second method for assessing binding site quality in addition to the Site_stats_rank_compare.pdf plot. Finally, R^2^ values for the residence times of each binding site are found within the Dataset.csv and BindingSites_Info_*Lipid*.txt files. These can also be used to assess whether CG simulations have been run for long enough to sufficiently sample protein-lipid interactions and to yield reliable outputs from PyLipID.
16. Exclude any poor binding sites from future analysis.

### Comparing lipid poses with cryo-EM densities

TIMING: Steps 17-19, 30 mins

17. Visualise the bound lipid poses outputted by PyLipID within the ‘*Interaction_Lipid/Bound_poses_Lipid*’ directory using VMD. Top ranked lipid binding poses (‘*BSidX_rank*’) and clustered poses (‘*BSidX_clusters*’) are located within subsidiary directories for each binding site.
18. Open software for visualisation of cryo-EM densities (e.g. Coot, Chimera, PyMOL) and load the protein coordinate file and corresponding density map.
19. Compare the identified binding poses with the position of densities in the cryo-EM structure. It is also possible to run LipIDens iterativly as e.g. map resolutions are improved and new sites become visible.

### Ranking site lipids

TIMING: Step 20, 1 min, Step 21, ~20 mins, Step 22, 1 min

20. In this stage of the protocol the residence times of different lipids binding to the same site are compared. Run the LipIDens python script and select the appropriate stage (‘5’). Binding site residues for different lipids are iteratively compared to those of the reference lipid (first lipid inputted). Sites which match most closely are selected across lipids species i.e. to compare different lipids binding to similar site locations. Corresponding binding sites IDs are written out in order and stored as a python dictionary BindingSite_ID_dict. Where corresponding sites could not be identified these are marked by ‘X’. Δ CRITICAL STEP: Check predicted binding site matches by comparing the lipid poses/binding sites from PyLipID. The BindingSite_ID_dict may need to be adjusted if a site was previously identified as poor (see ‘*Screening PyLipID data*’) or is assigned multiple times. If you are happy with predicted sites, accept the BindingSite_ID_dict, run the corresponding code and proceed to step 22. If not, follow step 21.
21. Within the notebook/master python script change the BindingSite_ID_dict dictionary keys to the lipids to compare. Set the corresponding values for each lipid to a list of binding site IDs for each corresponding site. For example, to compare the residence time of POPC binding site 2 and POPE binding site 4 (assuming these correspond to similar locations on the protein) then BindingSite_ID_dict={‘POPC’:[2], ‘POPE’: [4]}. If the lipid does not bind to a site then set the site ID to “X”.
Run the corresponding code to plot a comparison of residence times and R^2^ values for each site across lipid species. A ‘*Lipid_compare*’ directory containing plots (Lipid_compare_BSstats_PyLipID_Site_idx_*X_*ref*_Lipid*.pdf) for each site are generated where *X* corresponds to the reference lipid binding site ID number in BindingSite_ID_dict e.g. BindingSite_ID_dict={‘POPC’:[2, 3], ‘POPE’: [4, 5]} would produce two plots numbered Lipid_compare_BSstats_PyLipID_Site_idx_2_ref_POPC.pdf (comparing POPC site 2 with POPE site 4) and Lipid_compare_BSstats_PyLipID_Site_idx_3_ref_POPC.pdf (comparing POPC site 3 with POPE site 5) respectively.

i. The generated plots can be used to infer the most likely identity of a lipid species accounting for a density within the cryo-EM map (step 19).

### Lipid pose refinement using atomistic simulations

TIMING: Steps 23-24, 30 mins, Step 25, ~4-10 days (variable depending on available compute resources, simulation system size and length), Steps 26-27, 20 mins

23. The final section of the protocol is optional and relates to the refinement of CG lipid poses using atomistic simulations. Run the master python script and select the appropriate step (‘6’). Set the following user defined variables:

i. Set input_CG_frame to the CG simulation frame to use for back-mapping to atomistic resolution. Specified CG frames can be written to coordinate files using gmx trjconv with the -dump flag. The replicate and frame from which a lipid binding pose was obtained is noted within the pose_info.txt file (within with ‘*BSidX_rank/cluster*’ subdirectories).
ii. Set protein_AT_full to the atomistic structure used to establish CG simulations during the first stage of the protocol. To use an alternative input structure (e.g. including alternative protonation states or conformations), redefine protein_AT_full within this section as the modified input protein pdb file.
iii. Set model_type to either ‘de_novo’ or ‘aligned’ to select the output model from CG2AT^2^ to use for atomistic simulations. In the *de novo* model the protein coordinates are mapped to their positions within the CG frame. In the aligned model the protein coordinates are mapped to those of the input atomistic pdb (protein_AT_full).
iv. Change replicates_AT to the number of atomistic simulation replicates.
v. Change AT_simulation_time to the simulation time in nanoseconds.
24. Run the code corresponding to atomistic simulation setup (up to the stage marked ‘PAUSE POINT’). This is done automatically in the master python script. The output of this step is a GROMACS md.tpr file for each atomistic simulation replicate. An outline of the commands implemented within this section is given below:

i. Convert the protein-lipid system from CG to atomistic resolution using CG2AT, details of which are provided in ^2^. The protein conformation is backmapped either based on the coordinates in the CG frame (model_type=‘de_novo’) or those of the atomistic structure (model_type=‘aligned’). Lipid coordinates are backmapped to their positions in the CG frame. Users may select an atomistic forcefield and water model to build the system. The system is energy minimised and equilibrated. ?TROUBLESHOOTING
ii. Generate the .tpr file for the production run using gmx grompp.
25. Run the atomistic simulation using gmx mdrun. ?TROUBLESHOOTING. It is recommended to offload this calculation to a cluster or high-performance computing facility. □PAUSE POINT: wait until atomistic simulations have finished running before proceeding.
26. Once the atomistic simulations have finished running, compare the refined lipid binding pose to the cryo-EM density by loading in a preferred visualisation software.
27. Evaluate the match between the simulation derived lipid pose and the cryo-EM density using Q scores^23^. Details on implementation of this step in UCSF Chimera are given below:

i. Align the simulation frame with the atomistic input structure to which the density map corresponds. In PyMOL this can be done using the align or cealign commands, selecting the protein Cα or backbone beads of each structure (e.g. cealign *structure_name* and name CA, *simulation_frame_name* and name CA). It is often easier to remove superfluous components (i.e. everything except the protein and lipid of interest) from the system and save as a new .pdb file.
ii. Open UCSF Chimera and ensure the MapQ plugin^23^ is installed (for details see https://github.com/gregdp/mapq).
iii. Load the aligned simulation frame and cryo-EM map.
iv. Open the MapQ plugin using Tools > Volume Data > MapQ.
v. Enter the resolution of the cryo-EM map in the box marker ‘Res:’ and click ‘Calc’ to calculate Q scores.
vi. After the Q score calculation has finished. Select a protein sequence in proximity to the lipid using Ctrl-D and click to show the per atom Q scores on the structure. For further details see https://github.com/gregdp/mapq/tree/master/tutorials. Q scores are also mapped to the B-factor column of an output .pdb file from MapQ. It is expected that low/ negative Q scores may be observed for lipid regions outside of observed densities due to increased fluctuation of non-bound lipid regions.

## Troubleshooting

Troubleshooting advice can be found in Table 1.

**Table 1:** Troubleshooting table.

| Step | Problem | Possible Reason | Solution |
| --- | --- | --- | --- |
| 5 | Script does not produce required CG outputs | GROMACS or step errors<br>Incorrect input file locations<br>Excess protein segments<br>`insane.py` does not include a particular lipid | Step outputs are located within the `'output_files'` directory for each replicate. These can be used to diagnose setup failures.<br>Ensure correct paths to files defined<br>Try using the `-merge` flag in the `martinize` command<br>Add lipid to `insane.py` script |
| 4iv, 5ii | Protein embedded in membrane in incorrect orientation | Atomistic structure incorrectly orientated | Change the `protein_shift` and `protein_rotate` variables until satisfied with the orientation |
| 5v, 5vi, 6, 25 | Simulation crashes | Minimisation failed<br>Equilibration failed | Check atom resulting in failed minimisation to see if e.g. atom overlay results from a trapped water or lipid tail and remove if possible<br>Alter force constant and cut-off of the elastic network in the `martinize` command<br>Minimise atomistic structure before converting to CG resolution |
|  |  |  | Check .mdp file parameters |
| 24i | CG2AT fails | Lipid type not in database<br>CG frame not aligned within the PBC of the box resulting in incorrectly constructed system such as, e.g. water within the bilayer region or lipids out of the bilayer<br>Significant instability/clashes in input CG frame | Add the lipid to the database with corresponding atoms to map for the atomistic forcefield<br>Try extracting a CG frame from a centred trajectory or rotate the system using `gmx editconf`<br>Minimise the CG frame or select a different frame for input |

## Time Taken

Steps 1-3, Structure processing: ~20 mins

Steps 4-6, Setting up and performing coarse-grained simulations: ~1 week

Steps 7-10, Testing PyLipID cut-offs: ~4 h

Steps 11-12, Selecting PyLipID input parameters and running PyLipID analysis: ~3 h

Steps 13-16, Screening PyLipID data: 5 mins

Steps 17-19, Comparing lipid poses with cryo-EM densities: ~30 mins

Steps 20-22, Ranking site lipids: ~20 mins

Steps 23-27, Lipid pose refinement using atomistic simulations: ~1 week

## Associated publications

XXX – LipIDens manuscript DOI placeholder.

## Acknowledgements

The LipIDens pipeline was developed by T.B.A., W.S., R.A.C. and A.L.D. who contributed methodological development of PyLipID. Code testing was performed by R.A.C., C.K.C. and M.M.G.G. Development of the LipIDens pipeline was assisted by the expertise of C.E.C., L.C., T.R., C.S., A.B.W., P.J.S. and M.S.P.S. T.B.A, C.E.C and M.M.G.G. are supported by Wellcome (102164/Z/13/Z). W.S. was supported by a Newton International Fellowship. R.A.C., A.L.D. M.S.P.S. and P.J.S. are funded by Wellcome (208361/Z/17/Z). A.L.D. has been additionally supported by BBSRC (BB/R00126X/1) and the Department of Biochemistry. C.K.C. is funded by the BBSRC (BB/S003339/1). P.J.S. is also supported by the BBSRC (BB/P01948X/1, BB/R002517/1, and BB/S003339/1), and the MRC (MR/S009213/1) and M.S.P.S.’s research is supported by the BBSRC (BB/R00126X/1) and PRACE (Partnership for Advanced Computing in Europe, 2016163984). C.S. is supported by Cancer Research UK (C20724/A26752), the BBSRC (BB/T01508X/1) and the European Research Council (647278). L.C. is supported by a Wellcome administrative support grant (203141/Z/16/Z). A.B.W. acknowledges support from the Ray Thomas Edwards Foundation.

